# Reconstructing the Evolutionary History of Nitrogenases: Evidence for Ancestral Molybdenum-Cofactor Utilization

**DOI:** 10.1101/714469

**Authors:** Amanda K. Garcia, Hanon McShea, Bryan Kolaczkowski, Betül Kaçar

## Abstract

The nitrogenase metalloenzyme family, essential for supplying fixed nitrogen to the biosphere, is one of life’s key biogeochemical innovations. The three isozymes of nitrogenase differ in their metal dependence, each binding either a FeMo-, FeV-, or FeFe-cofactor where the reduction of dinitrogen takes place. The history of nitrogenase metal dependence has been of particular interest due to the possible implication that ancient marine metal availabilities have significantly constrained nitrogenase evolution over geologic time. Here, we reconstructed the evolutionary history of nitrogenases, and combined phylogenetic reconstruction, ancestral sequence inference, and structural homology modeling to evaluate the potential metal dependence of ancient nitrogenases. We find that active-site sequence features can reliably distinguish extant Mo-nitrogenases from V- and Fe-nitrogenases, and that inferred ancestral sequences at the deepest nodes of the phylogeny suggest these ancient proteins most resemble modern Mo-nitrogenases. Taxa representing early-branching nitrogenase lineages lack one or more biosynthetic *nifE* and *nifN* genes that both contribute to the assembly of the FeMo-cofactor in studied organisms, suggesting that early Mo-nitrogenases may have utilized an alternate and/or simplified pathway for cofactor biosynthesis. Our results underscore the profound impacts that protein-level innovations likely had on shaping global biogeochemical cycles throughout the Precambrian, in contrast to organism-level innovations that characterize the Phanerozoic Eon.

## 1 INTRODUCTION

All known life requires nitrogen for the synthesis of essential biomolecules, including nucleotides and amino acids. However, though the atmosphere contains nearly 80% N_2_ by volume, most organisms are not able to assimilate N_2_. Select bacteria and archaea called diazotrophs accomplish biological nitrogen fixation by nitrogenase metalloenzymes (E.C. 1.18.6.1) that catalyze the reduction of N_2_ to bioavailable NH_3_. Both phylogenetic and geochemical evidence suggest that nitrogenases are an ancient family of enzymes; time-calibrated phylogenies place the evolution of nitrogenases at ∼1.5–2.2 billion years ago (Ga) (Boyd, Anbar, et al., 2011), and the oldest potential isotopic biosignatures of nitrogenase activity date even further back to ∼3.2 Ga (Stueken, Buick, Guy, & Koehler, 2015). Because nitrogen has been suggested to be an important limiting nutrient on geologic timescales (Falkowski, 1997), nitrogenases have likely played a key role in the expansion of the biosphere for much of Earth’s history.

The nitrogenase family consists of three homologous isozymes (Boyd, Hamilton, & Peters, 2011; Raymond, Siefert, Staples, & Blankenship, 2004) named after the differential metal content of the active-site cofactor: Mo-nitrogenase (Nif, encoded by *nif*), V-nitrogenase (Vnf, encoded by *vnf*), and Fe-nitrogenase (Anf, encoded by *anf*) (Bulen & LeComte, 1966; Eady, 1996; Joerger & Bishop, 1988; Mus, Alleman, Pence, Seefeldt, & Peters, 2018) (Figure 1). Of the three isozymes, Mo-nitrogenases are the most common and widely studied; V- and Fe-nitrogenases are comparatively rarer and only known in taxa that also possess Mo-nitrogenase (Boyd, Hamilton, et al., 2011; Dos Santos, Fang, Mason, Setubal, & Dixon, 2012). All three nitrogenase isozymes are structurally and functionally similar, each containing two protein components: the electron delivery component (NifH, VnfH, or AnfH) is a homodimer and the catalytic component is either an α_2_β_2_ heterotetramer (MoFe protein, NifDK) or an α_2_β_2_γ_2_ heterohexamer (VFe protein, VnfDGK or FeFe protein, AnfDGK) (Figure 1a) (Bulen & LeComte, 1966; Hales, Case, Morningstar, Dzeda, & Mauterer, 1986; Schmid et al., 2002; D. Sippel & Einsle, 2017). During catalysis, the electron delivery component transiently associates with and delivers electrons to the catalytic component (Hageman & Burris, 1978). Electrons accumulate at the active site for N_2_ binding and reduction (Hoffman, Lukoyanov, Yang, Dean, & Seefeldt, 2014), which houses a homocitrate-metallocluster cofactor unique to each isozyme: the FeMo-cofactor in Nif (Figure 1b), FeV-cofactor in Vnf (Figure 1c), and FeFe-cofactor in Anf (Figure 1d) (Eady, 1996; Harris, Lukoyanov, et al., 2018; Krahn et al., 2002; Mus et al., 2018; D. Sippel & Einsle, 2017; Spatzal et al., 2011). Available spectral evidence suggests that these cofactors are structurally similar, with the main difference being the substitution of a Mo, V, or additional Fe atom (FeV-cofactor is also proposed to incorporate a carbonate ligand in place of one sulfur atom (D. Sippel & Einsle, 2017)) (Eady, 1996; Krahn et al., 2002; Spatzal et al., 2011). Nevertheless, biochemical studies demonstrate variable catalytic properties among the three nitrogenase isozymes, including differential abilities to reduce alternative substrates (Harris, Lukoyanov, et al., 2018; Harris, Yang, Dean, Seefeldt, & Hoffman, 2018; Hu et al., 2018; Hu, Lee, & Ribbe, 2011; Zheng et al., 2018). These catalytic variations likely arise due to a combination of the aforementioned cofactor compositional differences as well as differences in the surrounding protein environment (Fixen et al., 2016; Harris, Yang, et al., 2018; Lee et al., 2018; Rebelein, Lee, Newcomb, Hu, & Ribbe, 2018; Zheng et al., 2018).

**Figure 1.**
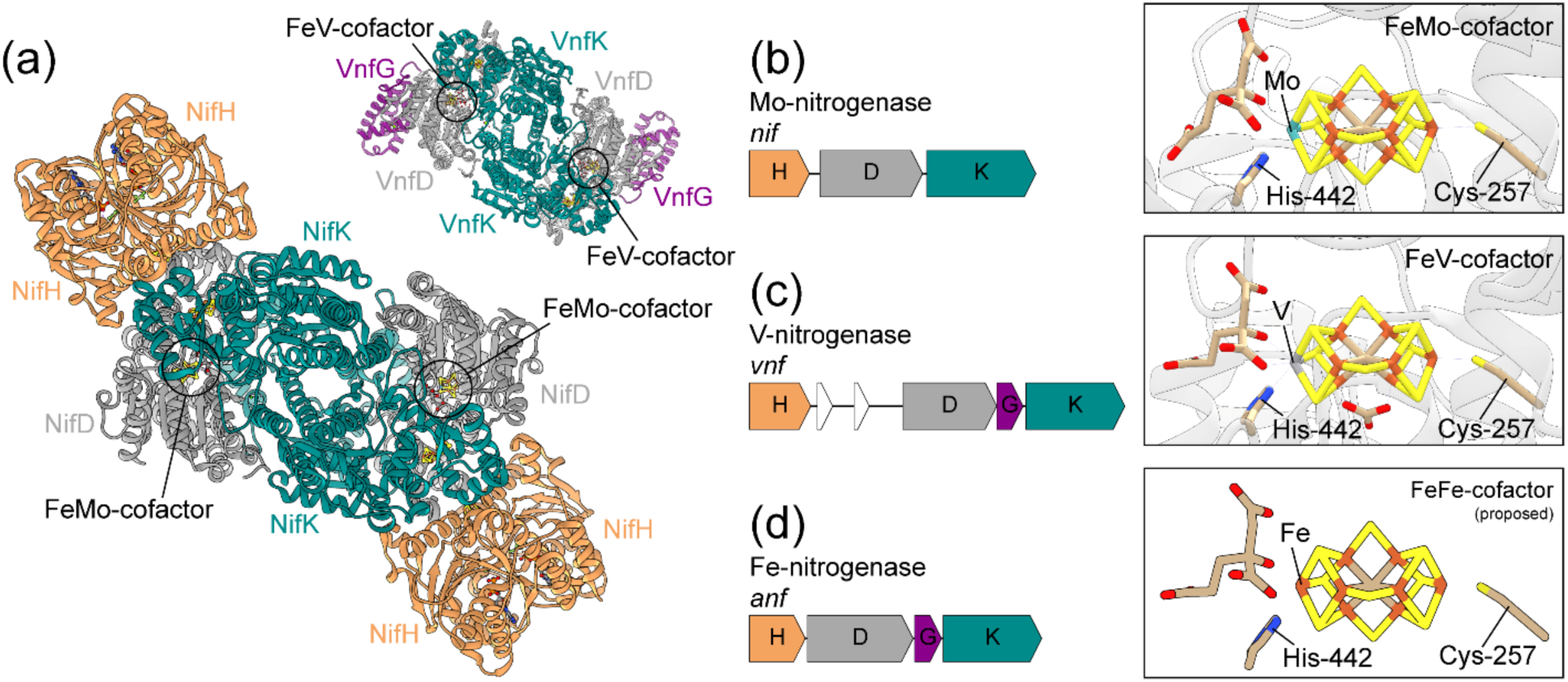
Structure and genetics of the three nitrogenase isozymes. (a) Structure of the *A. vinelandii* Mo-nitrogenase enzyme complex (NifHDK; PDB 1M34 (Schmid et al., 2002)) and V-nitrogenase VFe-protein component (VnfDGK; PDB 5N6Y (D. Sippel & Einsle, 2017)) (Fe-nitrogenase structure not previously published). The active-site FeMo-cofactor of Mo-nitrogenase and FeV-cofactor of V-nitrogenase are circled. (b-d) Catalytic genes and cofactor structures of *A. vinelandii* Mo-nitrogenase (b; PDB 3U7Q (Spatzal et al., 2011)), V-nitrogenase (c; PDB 5N6Y (D. Sippel & Einsle, 2017)), and Fe-nitrogenase (d; proposed structure (Harris, Lukoyanov, et al., 2018)). Residue numbering from aligned *A. vinelandii* NifD. Cofactor atom coloring is as follows: C, tan; Fe, rust; Mo, cyan; N, blue; O, red; S, yellow.

Metal cofactor incorporation in nitrogenases is constrained at multiple levels. At the level of single enzyme functionality, nitrogenase biochemical and biophysical properties shape metal-binding specificity. At a higher level, constraints arise from the partner proteins that constitute the biosynthetic mechanism for active-site cofactor assembly and insertion, best studied in Mo-nitrogenases (Curatti et al., 2007; Hu & Ribbe, 2011; Rubio & Ludden, 2008). In the *Azotobacter vinelandii* (*A. vinelandii*) model and *in vitro* systems, FeMo-cofactor assembly requires several associated proteins encoded within the *nif* gene cluster (Curatti et al., 2007; Hu, Fay, & Ribbe, 2005; Shah, Allen, Spangler, & Ludden, 1994; Shah, Imperial, Ugalde, Ludden, & Brill, 1986; Tal, Chun, Gavini, & Burgess, 1991). Most notably, the biosynthetic *nifB, nifE,* and *nifN* genes are considered, in addition to the catalytic *nifHDK* genes, to be — perhaps minimally — required for FeMo-cofactor assembly and Mo-nitrogenase function (Boyd, Anbar, et al., 2011; Curatti et al., 2007; Dos Santos et al., 2012; Hu et al., 2005; Shah et al., 1994; Shah et al., 1986; Tal et al., 1991). In *A. vinelandii, nifE* and *nifN* loci are located just downstream of the *nifHDK* cluster, whereas the *nifB* locus is located within a separate *nif* region near other regulatory and biosynthetic *nif* genes (Setubal et al., 2009). NifB catalyzes the formation of a Fe-S-C metallocluster, a precursor that forms the core of the mature FeMo-cofactor (Allen, Chatterjee, Ludden, & Shah, 1995; Hu & Ribbe, 2011). This precursor metallocluster is then transferred to a protein heterotetramer composed of NifE and NifN subunits (Allen et al., 1995; Roll, Shah, Dean, & Roberts, 1995), homologous to NifD and NifK, respectively, and likely having arisen by gene duplication (Boyd, Anbar, et al., 2011). Within NifEN, the precursor is further modified via the addition of homocitrate and Mo, and the mature cofactor is subsequently transferred to the nitrogenase NifDK catalytic protein component (Roll et al., 1995). Unlike that for the FeMo-cofactor, the biosynthetic pathways for the formation of the FeV- and FeFe-cofactors are relatively unknown. However, transcriptional profiling of the three nitrogenase systems in *A. vinelandii* suggests that FeV- and FeFe-cofactor synthesis relies on several *nif* genes in addition to *vnf* and *anf* genes, respectively (Hamilton et al., 2011; Joerger & Bishop, 1988; Kennedy & Dean, 1992). These include *nifBEN* for most alternative gene clusters, with the exception of certain taxa (including *A. vinelandii*) that possess *vnfEN* homologs of *nifEN* that likely perform a similar biosynthetic function (Boyd, Anbar, et al., 2011; Boyd & Peters, 2013; Hamilton et al., 2011).

Paleobiological interest in nitrogenases has primarily centered on the coevolution of nitrogenase metal usage and the geochemical environment, with the possible implication that marine metal availabilities have significantly constrained nitrogenase evolution over geologic time (Anbar & Knoll, 2002; Boyd, Hamilton, et al., 2011; Canfield, Glazer, & Falkowski, 2010; Raymond et al., 2004). Inferences of ancient nitrogenase metal usage have relied on isotopic biosignatures (Stueken et al., 2015) and metal abundances (Anbar & Knoll, 2002) evidenced by the geologic record, as well as on phylogenetic reconstructions of both catalytic and cofactor biosynthesis proteins (Boyd, Anbar, et al., 2011; Boyd, Hamilton, et al., 2011; Raymond et al., 2004). High marine Fe concentrations and potential Mo scarcity prior to increased atmospheric oxygenation surrounding the ∼2.3-2.5 Ga Great Oxidation Event (Anbar et al., 2007; Lyons, Reinhard, & Planavsky, 2014) has led to the hypothesis that Fe- or V-nitrogenases may have been dominant in early oceans (Anbar & Knoll, 2002; Canfield et al., 2010) and possibly predate Mo-nitrogenases (Raymond et al., 2004). More recent phylogenetic reconstructions have instead suggested that the evolution of Mo-nitrogenases, dated by time-calibrated phylogenies of Nif/Vnf/AnfDKEN sequences to ∼1.5–2.2 Ga (Boyd, Anbar, et al., 2011), preceded that of V- and Fe-nitrogenases (Boyd, Hamilton, et al., 2011). These phylogenetic inferences are also consistent with the observation that *vnf* and *anf* genes are only present in organisms that also harbor *nif*, and that V-/Fe-nitrogenase assembly relies on *nif* biosynthetic genes (Hamilton et al., 2011; Joerger & Bishop, 1988; Kennedy & Dean, 1992). However, ∼3.2-Ga isotopic signatures of biological nitrogen fixation suggest an earlier origin of nitrogenase (Stueken et al., 2015), and, even though isotopically consistent with Mo-dependent nitrogen fixation, predate age estimates of both Mo-nitrogenase (Boyd, Anbar, et al., 2011) and earliest marine Mo availability (Anbar et al., 2007; Anbar & Knoll, 2002; Lyons et al., 2014). Thus, the evolutionary trajectory of nitrogenase metal usage — and by extension the link between nitrogenase evolution and marine metal availabilities over geologic time — is not yet known.

Here, we explored the indicators of nitrogenase metal usage history by a combined method relying on ancestral sequence reconstruction, an evolutionary approach by which inferred, historical protein sequence information can be linked to functional inferences of molecular properties evidenced by computational analyses or laboratory experiments (Aadland, Pugh, & Kolaczkowski, 2019; Benner, Sassi, & Gaucher, 2007; Thornton, 2004). These paleogenetic approaches have been increasingly applied in biogeochemically relevant molecular studies to offer insights into the coevolution of life and Earth (Garcia & Kacar, 2019; Gomez-Fernandez et al., 2018; Kacar, Hanson-Smith, Adam, & Boekelheide, 2017). We reconstructed the phylogenetic history of Mo-, V-, and Fe-nitrogenases in order to resurrect ancestral nitrogenases *in silico*, as well as to map the taxonomic distribution of cofactor biosynthetic components considered necessary for cofactor assembly (Boyd, Anbar, et al., 2011; Curatti et al., 2007; Dos Santos et al., 2012; Hu et al., 2005; Shah et al., 1994; Shah et al., 1986; Tal et al., 1991). Through this combined approach, we access ancestral enzyme sequence features suggestive of ancient Mo-dependence. We further present phylogenetic data that suggests that the ancient Mo-utilization was potentially achieved via a unique pathway for the synthesis of the FeMo- or similar cofactor that may be present in poorly characterized, basal lineages. Integration of protein evolution and paleobiology is a unique melding of disparate data sets; this approach may provide hypotheses that address interactions ranging from the external environment to the cellular environment, and from the cellular environment to that maintained around the interacting protein. The exchange of materials across these different scales necessitates constraints on the flow and availability of substrates that make such exchanges possible. It is the specific nature of these constraints and how they may change in response to external perturbations that enable us to develop completely new experimentally testable hypotheses that connect geochemical reservoirs with biological metabolisms — those that cannot be constructed from macroevolutionary or geological frameworks alone.

## 2 MATERIALS AND METHODS

### 2.1 Ancestral reconstruction of nitrogenase protein sequences

An initial dataset of extant nitrogenase Nif /Vnf/AnfHDK homologs was constructed by retrieving amino acid sequences from the National Center for Biotechnology Information non-redundant protein database, accessed September 2018 (O’Leary et al., 2016). Potential homologs were identified by BLASTp (Camacho et al., 2009) using query sequences from *A. vinelandii* (NifH: Avin_01380, NifD: Avin_01390, NifK: Avin_01400) and an Expect value cutoff of <1e-5. The dataset was then manually curated to remove partial and distantly related sequences. Additional nitrogenase sequences were manually retrieved from the Joint Genome Institute Integrated Microbial Genomes and Microbiomes database, accessed September 2018 (Chen et al., 2019). The nitrogenase sequence dataset was finalized to include NifHDK sequences from 256 taxa, AnfHDK sequences from 14 taxa, VnfHDK sequences from 14 taxa, and outgroup light-independent protochlorophyllide oxidoreductase (Bch/ChlLNB) sequences — sharing distant homology with nitrogenases (Boyd, Anbar, et al., 2011; Hu & Ribbe, 2015; Raymond et al., 2004) — from 10 taxa (Appendix S1; additional analyses were performed with an expanded outgroup, Appendix S2). Only one Nif/Anf/VnfHDK sequence set was retained per genus to broaden taxonomic sampling. Equal sequence sampling for Anf and Vnf was made to remove the potential for oversampling bias in ancestral sequence inference. H-, D-, and K-subunit sequences corresponding to each taxon were manually checked for synteny of their encoding genes. AnfHDK and VnfHDK sequences were identified by the proximity of each gene locus to *anfG* or *vnfG*, which encodes the additional G-subunit present in the VeFe or FeFe protein, respectively, but not present in the MoFe protein (Eady, 1996). Finally, the presence of the cofactor biosynthetic *nifBEN* genes was investigated for all taxa represented in our dataset by BLASTp, as well as by manually inspecting the *nif* genome region.

Reconstruction of ancestral nitrogenase sequences was performed by PhyloBot (Hanson-Smith & Johnson, 2016) (www.phylobot.com), which automates multiple sequence alignment, phylogenetic reconstruction, and ancestral sequence inference methods. The concatenated 294-sequence dataset of Nif/Anf/VnfHDK homologs (including 10 Bch/ChlLNB outgroup sequences) was aligned by MSAProbs v0.9 5r1 (Liu, Schmidt, & Maskell, 2010) and MUSCLE v3.8.31 (Edgar, 2004). Both alignment outputs were then used to perform phylogenetic reconstruction by RAxML v8.1.15 (Stamatakis, 2014) under 6 different combinations of amino acid substitution and rate heterogeneity models. Branch support was evaluated by the approximate likelihood ratio test (aLRT) (Anisimova & Gascuel, 2006), which assesses the gain in overall likelihood against a null hypothesis of branch length = 0. Additional phylogenetic reconstructions with an expanded outgroup were performed outside of Phylobot to resolve root positioning, but were not used in subsequent ancestral sequence inference (Appendix S2). Ancestral sequences were inferred by joint maximum likelihood using CODEML v4.2 (Yang, 2007) at all nodes within the 12 Phylobot-constructed phylogenies (Tree-1 – Tree-12), with gaps inferred by parsimony. To assess ancestral sequence robustness to phylogenetic uncertainty (Hanson-Smith, Kolaczkowski, & Thornton, 2010), ancestors inferred from the top five phylogenies ranked by log likelihood scores were selected for further analysis (Table 1, Appendix S2). Finally, to evaluate the effects of ambiguously reconstructed sites on subsequent structural analyses, Bayesian sampled ancestors were inferred from the maximum likelihood site posterior probabilities calculated by CODEML (Aadland et al., 2019). 100 random Bayesian sequences were generated for each of five ancestral nodes of interest across the top five phylogenies. Thus, 25 maximum likelihood and 2,500 Bayesian-sampled ancestral sequences were analyzed in total. All maximum likelihood reconstructed trees and ancestral sequences are available for view and download at http://phylobot.com/613282215/.

**Table 1.**
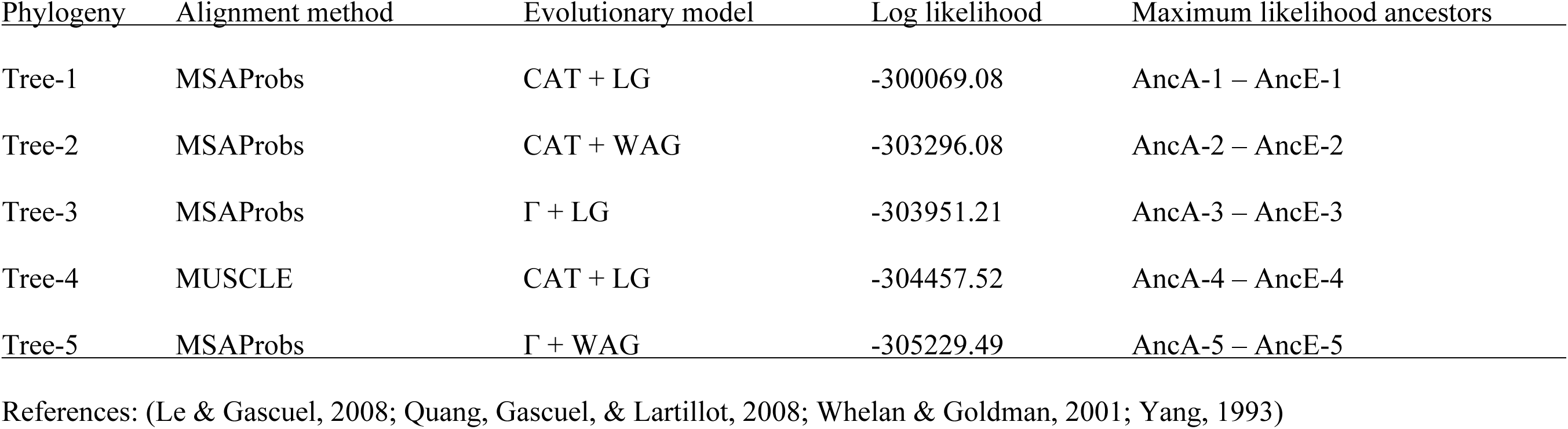
Alignment and evolutionary model parameter combinations for the top five phylogenies, ranked by log likelihood scores.

### 2.2 Probabilistic model for metal-dependence classification of nitrogenase sequences

Extant nitrogenase sequences of known metal dependence were labelled as binding either the FeMo- (“Nif”), FeV- (“Vnf”), or FeFe-cofactor (“Anf”) according to their phylogenetic clustering, and were used to train a support vector classifier that identifies cofactor specificity, based on the residues in the active-site cofactor-binding pocket. The 30 active-site residues were identified as those residing within 5 Å of any atom in either the FeMo-cofactor of the *A. vinelandii* NifD protein (PDB 3U7Q (Spatzal et al., 2011)) or the FeV-cofactor of the *A. vinelandii* VnfD protein (PDB 5N6Y (D. Sippel & Einsle, 2017)) (see Section 3.3 for further discussion of selected residues). The corresponding residues in other extant homologs and inferred ancestral sequences were identified by multiple sequence alignment using MAFFT (Katoh & Standley, 2013). For each sequence in the labelled dataset and residue in the active-site, the probability distribution over all 20 possible amino acids was inferred by averaging (1) the observed distribution for that sequence (which places probability 1.0 on the observed residue, (2) the probability distribution that would be expected over a short evolutionary distance (0.01 substitutions/site), starting from the observed residue and assuming the JTT substitution model (Jones, Taylor, & Thornton, 1992), and (3) a diffuse Dirichlet mixture regularizer that places relatively low probabilities across all possible amino acids (Sjolander et al., 1996). The labelled dataset was randomly divided into 60% training data for training the support vector classifier and 40% out-of-sample testing. The support vector classifier used a radial basis kernel, with the kernel coefficient estimated as 1 / (# of predicted features × Var(features)). Data samples were weighted based on the frequencies of Nif, Vnf, and Anf labels in the training data: the weight assigned to samples labelled as cofactor *I* (c_i_) = # of samples / (3 × Freq(c_i_)). The regularization parameter was optimized using 5-fold cross-validation, and classification accuracy was assessed by out-of-sample testing. The procedure of randomly splitting data into training/testing, training the classifier, and assessing out-of-sample accuracy was replicated 10 times.

Extant sequences and maximum-likelihood ancestral-reconstructed sequences were classified as Nif, Vnf, or Anf using one-vs-rest multi-class classification, based on the observed amino-acid residues in their active sites. In addition, ancestral sequences were also classified based on the inferred posterior-probability distributions over all 20 possible amino acids at each position in the active site. Classification support is reported as the distance of each data sample from the support-vector hyperplane defining the inferred class.

### 2.3 Structural homology modeling of extant and ancestral nitrogenase D-subunits

Structural homology modeling of extant and ancestral (25 maximum likelihood and 2,500 Bayesian-sampled) nitrogenase D-subunit proteins was performed by Modeller v9.2 (Sali & Blundell, 1993). Extant nitrogenase sequences, broadly sampled from the reconstructed nitrogenase phylogeny, were modeled to provide comparisons with ancestral models. D-subunit sequences from extant and ancestral nitrogenases were aligned to 38 NifD and 2 VnfD structural templates retrieved from the Protein Data Bank (Berman et al., 2000), accessed November 2018 (Appendix S3; published AnfD models not available at time of analysis). Information from all 40 templates was used to model each structure. All models were generated by specifying the inclusion of the FeMo-cofactor of the 3U7Q NifD structure (Spatzal et al., 2011), selected as the highest resolution Mo-nitrogenase template. To assess the effect of the template cofactor type on the generated structure, additional models were constructed by specifying the inclusion of the FeV-cofactor of the 56NY VnfD template (D. Sippel & Einsle, 2017) (Appendix S3). Modeling replicates (100 in total) were performed per sequence and assessed by averaging over the scaled Modeller objective function, Discrete Optimized Protein Energy, and high resolution Discrete Optimized Protein Energy scores, as previously described (Aadland et al., 2019). The ten best modeling replicates per extant sequence, ten best replicates per maximum likelihood ancestral sequence, and the single best replicate per Bayesian-sampled variant sequence were selected for further analysis, totaling 3,080 models with the FeMo-cofactor specified.

### 2.4 Active-site pocket volume calculation of extant and ancestral D-subunit models

Volumes of the modeled ancestral and extant D-subunit active-site cofactor pockets were calculated by POVME v2.0 (Durrant, Votapka, Sorensen, & Amaro, 2014). Spatial coordinates and the inclusion region for volume calculation were specified manually. Pocket volumes were calculated with a grid spacing of 0.5 Å and a 1.09 Å distance cutoff from any receptor atom’s van der Waals radius. Volume outside of the modeled convex hull of the cofactor pocket as well as noncontiguous volume were removed. Statistical analysis of ancestral and extant pocket volume data was performed in R (R Core Team, 2014).

## 3 RESULTS

### 3.1 V- and Fe-nitrogenases diversified after Mo-nitrogenases

We reconstructed the phylogenetic history of Mo-, V-, and Fe-nitrogenases to infer ancestral nitrogenase sequences and associated indicators of nitrogenase metal dependence. Nif/Anf/VnfHDK protein homologs curated from the National Center for Biotechnology Information and Joint Genome Institute databases represent 20 bacterial and archaeal phyla, 11 of which are known from experimental investigations to include diazotrophic taxa (Dos Santos et al., 2012; Ormeño-Orrillo, Hungria, & Martinez-Romero, 2013) (Appendix S1). The five most represented phyla in our dataset — Bacteroidetes, Cyanobacteria, Firmicutes, Proteobacteria, and Euryarchaeota — encompass ∼80% of the curated sequences. Our genomic dataset also suggests the presence of nitrogen fixation within the Acidobacteria, Actinobacteria, Aquificae, Chlorobi, Chloroflexi, Chrysiogenetes, Deferribacteres, Elusimicrobia, Fusobacteria, Lentisphaerae, *Candidatus* Margulisbacteria, Nitrospirae, Planctomycetes, Spirochaetes, and Verrucomicrobia clades.

Our maximum likelihood nitrogenase phylogeny segregates Nif/Anf/VnfHDK sequences into two major lineages similarly observed in previous studies (Boyd, Anbar, et al., 2011; Raymond et al., 2004) (Tree-1; Figure 2): the first comprises Nif-I and Nif-II Mo-nitrogenases (primarily aerobic/facultative bacteria and anaerobic/facultative bacteria/archaea, respectively; highlighted in blue) (Boyd, Costas, Hamilton, Mus, & Peters, 2015; Raymond et al., 2004), and the second comprises V- and Fe-only-nitrogenases (Vnf and Anf, respectively; highlighted in red) as well as three clades of “uncharacterized” nitrogenases that lack extensive experimental characterization with regard to metal dependence (Mb-Mc, highlighted in purple; F-Mc, highlighted in green; Clfx, highlighted in yellow) (Boyd, Hamilton, et al., 2011; Dekas, Poretsky, & Orphan, 2009; Dos Santos et al., 2012; Mehta & Baross, 2006). We additionally analyzed four alternate phylogenies (ranked by log likelihood scores), reconstructed by varying alignment methods, amino acid substitution models, and rate heterogeneity models (Tree-2–Tree-5; Appendix S2). The aforementioned major nitrogenase clades are present across all alternate phylogenetic topologies. Branch ordering of these major clades is similarly consistent, with the exception of root position: in Tree-3 and Tree-5, the root is placed between Nif-I and Nif-II/uncharacterized/Anf/Vnf, rather than between Nif-I/Nif-II and uncharacterized/Anf/Vnf as in Tree-1, Tree-2, and Tree-4. Nevertheless, additional phylogenetic analyses incorporating an expanded outgroup provide stronger support for the root position of Tree-1, Tree-2, and Tree-4 (Appendix S2).

**Figure 2.**
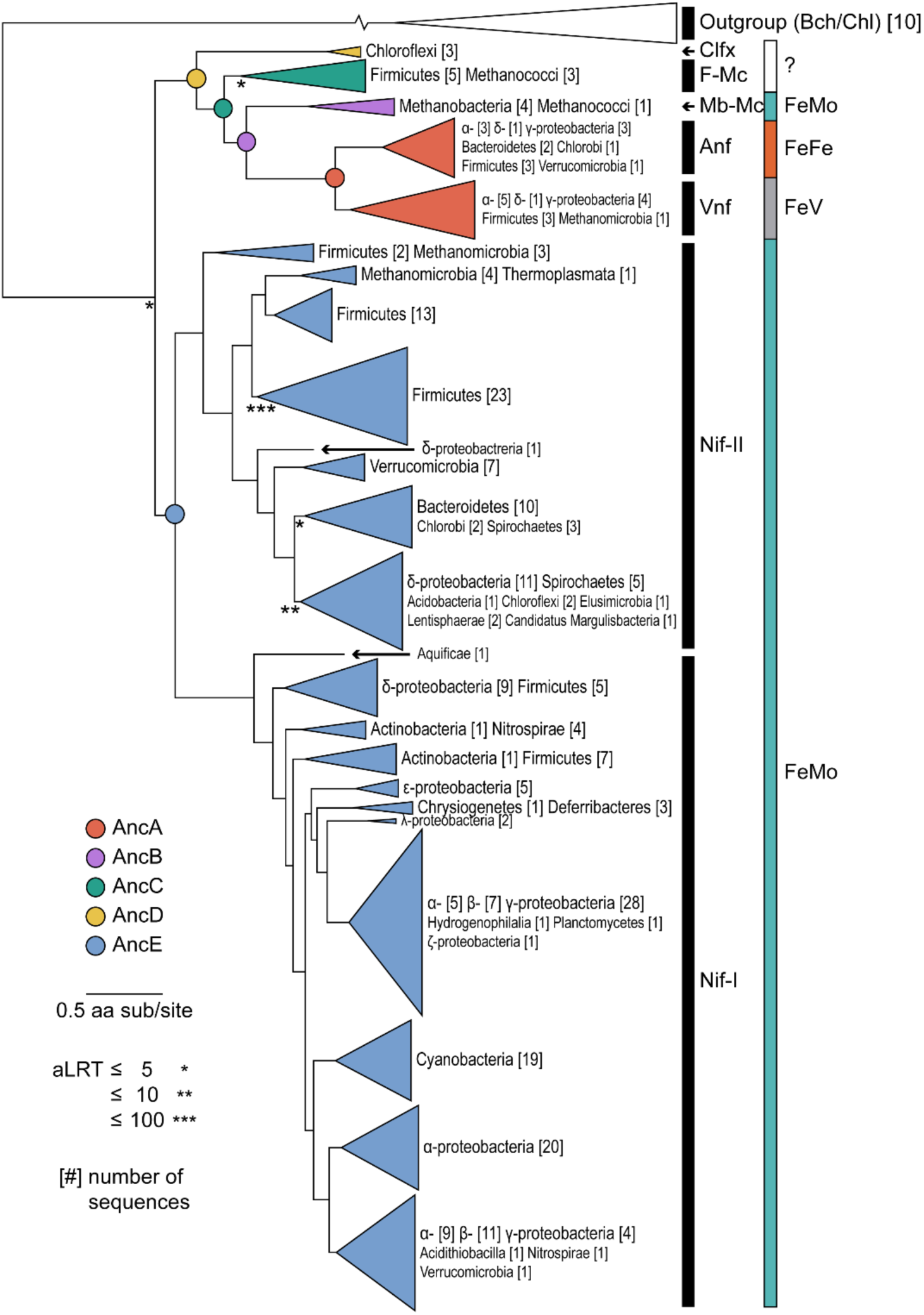
Maximum likelihood phylogeny of concatenated Nif/Anf/VnfHDK nitrogenase and Bch/ChlLNB outgroup protein sequences (Tree-1; see Table 1). Ancestral nodes analyzed in this study are labeled AncA–AncE. Known active-site cofactor metal content is listed on the right. Branch support is derived from the approximate likelihood ratio test (aLRT). Branch length scale is in units of amino acid substitutions per site. Outgroup branch break used to conserve space; true branch length = 5.578 substitutions per site. Phylogeny coloring is as follows: Clfx, yellow; F-Mc, green; Mb-Mc, purple; Anf/Vnf, red; Nif-I/-II, blue.

In all analyzed nitrogenase phylogenies, Vnf and Anf sequences (highlighted in red) form reciprocally monophyletic clades that branch immediately distal to Mb-Mc nitrogenases (highlighted in purple) (Figure 2, Appendix S2). The reciprocal monophyly of Vnf and Anf, as well as the clustering of Mb-Mc, Vnf, and Anf, is well-supported across all phylogenetic topologies (aLRT > 10^6^). The Mb-Mc clade is composed of archaeal hydrogenotrophic methanogens within classes Methanobacteria and Methanococci. Because all Mb-Mc taxa possess *nifBEN* genes considered to be minimally required for the synthesis of the FeMo-cofactor (Boyd, Anbar, et al., 2011; Curatti et al., 2007; Dos Santos et al., 2012; Hu et al., 2005; Shah et al., 1994; Shah et al., 1986; Tal et al., 1991), it is likely that Mb-Mc nitrogenases are Mo-dependent (Boyd, Hamilton, et al., 2011). Thus, the phylogenetic positioning of Vnf and Anf is consistent with previous suggestions that V- and Fe-nitrogenases diversified after Mo-nitrogenases (Boyd, Hamilton, et al., 2011).

### 3.2 Basal uncharacterized nitrogenases lack associated genes for FeMo-cofactor synthesis

In addition to investigating the phylogenetic relationships between Mo-, V-, and Fe-nitrogenase isozymes, we mapped the presence of the biosynthetic *nifB*, *nifE*, and *nifN* genes — considered to be necessary for FeMo-cofactor assembly (Boyd, Anbar, et al., 2011; Curatti et al., 2007; Dos Santos et al., 2012; Hu et al., 2005; Shah et al., 1994; Shah et al., 1986; Tal et al., 1991) — among taxa represented in our dataset. All analyzed taxa possess the full complement of *nifBEN* biosynthetic genes, with the exception of two uncharacterized clades: Clfx (highlighted in yellow) and F-Mc (highlighted in green) (Figure 2). Within the lineage containing Vnf and Anf nitrogenases, Clfx and F-Mc clades are most basal. These branching positions of Clfx and F-Mc clades within the Vnf/Anf lineage are consistently observed across all phylogenetic topologies (Trees-1–5; Appendix S2) and are well-supported (aLRT > 10^3^ for Clfx, aLRT > 10^18^ for F-Mc). In Tree-1-, 2-, and -4, as well as in trees reconstructed with an expanded outgroup (Appendix S2), Clfx and F-Mc clades also branch immediately distal to the root.

Both basal Clfx and F-Mc homologs are found primarily in thermophilic taxa and lack one or more associated *nifEN* cofactor biosynthesis genes, as has been noted previously (Boyd, Hamilton, et al., 2011; Dos Santos et al., 2012; Shock & Boyd, 2015). Clfx homologs, constituting the most basal uncharacterized clade within the uncharacterized/Vnf/Anf lineage, represent in the present dataset three mesophilic or thermophilic Chloroflexi species (Figure 2, highlighted in yellow). The *nif* clusters of Clfx taxa lack both *nifE* and *nifN* genes, and the typically continuous *nifHDK* genes observed in other taxa are instead interrupted by *nifB* (arranged *nifHBDK*) (Setubal et al., 2009). F-Mc homologs branch immediately distal to the Clfx clade and represent eight thermophilic Firmicutes and archaeal methanogen (class Methanococci) taxa (Figure 2, highlighted in green). F-Mc species possess biosynthetic *nifB* and *nifE* genes, but not *nifN*, with the exception of *Methanothermococcus thermolithotrophicus* that retains *nifN*. A previous study found that sequence features and modeled structural features of F-Mc nitrogenases closely resemble those of Mo-nitrogenases (McGlynn, Boyd, Peters, & Orphan, 2012). However, the absence of *nifN* in most F-Mc taxa suggests that these strains may not be capable of synthesizing the FeMo-cofactor, given previous *in vitro* and *in vivo* experiments demonstrating the requirement of both *nifEN* for FeMo-cofactor biosynthesis (Curatti et al., 2007; Hu et al., 2005; Shah et al., 1994; Shah et al., 1986; Tal et al., 1991). The lack of one or more *nifEN* genes in Clfx and F-Mc taxa might additionally suggest that such strains cannot express functional nitrogenases. Nitrogen fixation among Clfx taxa has not been definitively demonstrated; the Clfx species *Oscillochloris trichoides* has only been reported to reduce acetylene, an alternate substrate of nitrogenase (Keppen, Baulina, & Kondratieva, 1994; Keppen, Lebedeva, Troshina, & Rodionov, 1989). However, the F-Mc species *Methanocaldococcus sp.* FS406 (lacking *nifN*), as well as an uncharacterized, anaerobic methane-oxidizing archaeon not included in the present study, have been shown to assimilate isotopically labeled nitrogen (Dekas et al., 2009; Mehta & Baross, 2006). The ability of early-branching F-Mc nitrogenases to fix nitrogen in the absence of the full complement of *nifEN* genes may indicate an alternative and/or simplified pathway for cofactor assembly not used for other Mo-, V-, and Fe-nitrogenases.

### 3.3 High statistical support for ancestral nitrogenase active-site residues

We inferred ancestral sequences for each of the H-, D-, and K-subunits that constitute the nitrogenase enzyme complex (Figure 1) across five phylogenetic topologies (Tree-1–5; Appendix S2). Ancestral nitrogenase sequences were inferred for five well-supported internal nodes along a phylogenetic transect between Mo- (highlighted in blue) and V-/Fe-nitrogenases (highlighted in red) (Figure 2). The five targeted nodes are: AncA (ancestral to Anf and Vnf), AncB (ancestral to AncA and Mb-Mc), AncC (ancestral to AncB and F-Mc), AncD (ancestral to AncC and Clfx), and AncE (ancestral to Nif-I and Nif-II). Thus, AncA–D are nested, whereas AncE lies along a divergent lineage toward Nif-I and Nif-II Mo-nitrogenases. For further analyses, we selected the maximum likelihood ancestral sequence per target node from each of the five phylogenies (Tree-1–5), totaling 25 sequences. Due to differences in root position, identical AncE nodes were not present across all topologies and analogous nodes were instead selected (Appendix S2). Ancestral sequences are hereafter labeled with the tree likelihood rank from which they were inferred (e.g., AncA from Tree-1 is labeled AncA-1). All tree and ancestral sequence information can be found at http://phylobot.com/613282215/.

Mean site posterior probabilities for ancestral nitrogenase HDK sequences across all phylogenies range between ∼0.83 and 0.91, and for the highest-likelihood phylogeny (Tree-1), between ∼0.84 and 0.90 (Appendix S4). Ancestral sequence support generally decreases with increasing phylogenetic node age. For example, within the uncharacterized/V-/Fe-nitrogenase linage, AncA-1 has the highest mean posterior probability (0.90 ± 0.18) and AncD-1 has the lowest mean posterior probability (0.84 ± 0.22). Mean ancestral sequence probability for each node also does not deviate by more than ∼0.02 across each of the five phylogenetic topologies (Tree-1–5; Appendix S4). These observations suggest that sequence support for ancestral nitrogenases is more sensitive to ancestral node position than to topological differences between the analyzed trees.

In addition to surveying total ancestral HDK sequence support, we analyzed support for 30 active-site residues, defined as those residing within 5 Å of any atom in either the FeMo-cofactor of the *A. vinelandii* NifD protein (PDB 3U7Q (Spatzal et al., 2011)) or the FeV-cofactor of the *A. vinelandii* VnfD protein (PDB 5N6Y (D. Sippel & Einsle, 2017)) (Figure 3). These active-site residues are not contiguous but are instead scattered throughout the D-subunit sequence. Mean posterior probabilities of ancestral active-site residues, which range between 0.92 to 0.98 across all phylogenies, are consistently greater than those of entire reconstructed nitrogenase HDK sequences (0.83–0.91) (Appendix S4). Of the 30 active-site residues, only five sites have, in one or more ancestral sequences, plausible alternative reconstructions with posterior probabilities >0.30: sites 59, 69, 358, 360, 425, 441 (site numbering both here and hereafter based on *A. vinelandii* NifD). No ancestral sequences have more than three active-site residues with such plausible alternative reconstructions. Ten active-site residues are conserved across all analyzed extant nitrogenases: Val-70, Gln-191, His-195, Cys-275, Arg-277, Ser-278, Gly-356, Phe-381, Gly-424, His-442. These conserved residues are thus reconstructed in all ancestral nitrogenases unambiguously (site posterior probability = 1.00). Statistical support for ancestral active-site residues (greater than 0.92) underpins subsequent analyses of ancestral active-site properties that may inform inferences of nitrogenase metal dependence.

**Figure 3.**
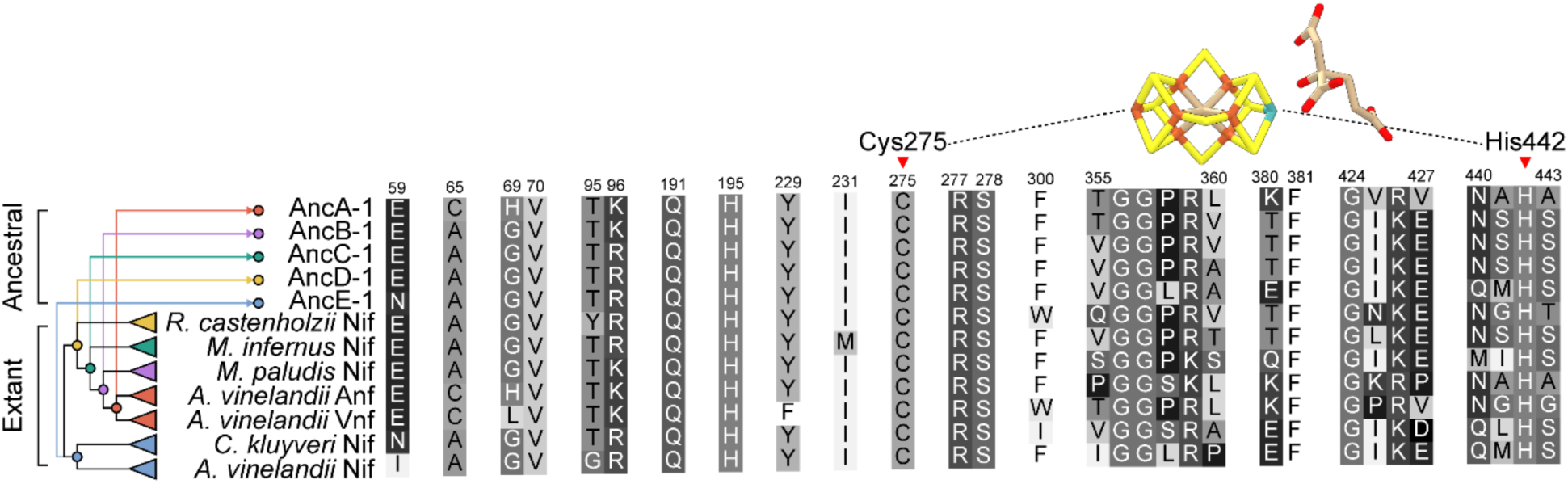
Active-site protein environment of representative ancestral and extant nitrogenases. All residues located within 5 Å of any atom in either the FeMo-cofactor of the *A. vinelandii* NifD protein (PDB 3U7Q (Spatzal et al., 2011)) or the FeV-cofactor of the *A. vinelandii* VnfD protein (5N6Y (D. Sippel & Einsle, 2017)). Residue numbering from aligned *A. vinelandii* NifD. Cys-275 and His-442 residues that coordinate the cofactor are indicated by red arrows. Phylogeny coloring is as follows: Clfx, yellow; F-Mc, green; Mb-Mc, purple; Anf/Vnf, red; Nif-I/-II, blue.

### 3.4 Oldest ancestral nitrogenase active-site sequences resemble extant Mo-nitrogenases

We analyzed sequence features of both ancestral and extant nitrogenases to identify those correlated with metal dependence. In particular, we focused on 30 nitrogenase active-site residues (Figure 3; see Section 3.3. for further details) for three reasons: (1) active-site residues are known to affect catalytic efficiency and substrate specificity (Brigle et al., 1987; Christiansen, Cash, Seefeldt, & Dean, 2000; Fixen et al., 2016; Kim, Newton, & Dean, 1995; Sarma et al., 2010; Yang, Moure, Dean, & Seefeldt, 2012) and thus may be tuned to nitrogenase metal dependence (2) active-site sequence features have previously been used, in part, to classify the metal dependence of extant uncharacterized nitrogenases (McGlynn et al., 2012) (3) statistical support for ancestral active-site residues are higher than the mean support across entire HDK sequences (see Section 3.3).

We first assessed the sensitivity of ancestral sequence variation to phylogenetic uncertainty and ancestral statistical support. Overall, mean identities for ancestral sequences compared across different nodes range from ∼55 to 90%. Ancestral sequences inferred from the same node across alternate phylogenies (Tree-1–5) have relatively high mean identities, ranging from ∼93 to 95% across the total HDK sequence and from ∼96 to 100% within the active site. These high mean identities suggest that topological differences among the alternate phylogenies used for ancestral sequence inference do not contribute to a high degree of ancestral sequence variation. Identities among sequences inferred from the same node also do not appear to be correlated with statistical support. For example, though full AncA HDK sequences are reconstructed with the highest mean statistical support (∼0.89–0.91), they exhibit neither the lowest nor the highest mean identities as compared with sequences inferred from other nodes (Appendix S4).

We next identified specific residues within the nitrogenase active site that are unique to particular isozymes of known metal dependence in order to survey their occurrence in ancestral sequences. We found that three active-site residues are unique to Mo-nitrogenases (Ala-65, Arg-96, Gln-440), six are unique to V-nitrogenases (Leu-69, Trp-300, Thr-355, Pro-358, Pro-425, Val-427), five are unique to Fe-nitrogenases (His-69, Pro-355, Lys-359, Pro-427, Ala-441), and six are unique to V- and Fe-nitrogenases (Cys-65, Lys-96, Leu-360, Lys-380, Arg-426, Asn-440) (Appendix S5). Though we are unaware of any experimental mutagenesis studies identifying a role for these uniquely conserved residues, we hypothesize that these residues may contribute to the metal specificity of the active-site environment. Surprisingly, most nitrogenase ancestors exhibit comparable numbers of residues unique to either V-/Fe-nitrogenases or Mo-nitrogenases, and thus their occurrence does not appear informative for inferring ancestral metal dependence (e.g., ancestral AncC sequences contain two residues unique to V-/Fe-nitrogenases and two residues unique to Mo-nitrogenases). An exception is AncA sequences, which contain a preponderance of active-site residues that are unique to extant V- and Fe-nitrogenases. One of the residues unique to only V-nitrogenases, Thr-355, has recently been suggested to interact directly with a proposed FeV-cofactor carbonate ligand not present in the FeMo-cofactor (D. Sippel & Einsle, 2017). This carbonate ligand lies within a secondary-structure loop that also contains Pro-358, unique to V-nitrogenases, and Leu-360, unique to V- and Fe-nitrogenases. These Thr-355, Pro-358, and Leu-360 residues are observed across all AncA sequences.

In addition to analyzing specific active-site residues, we compared the total active-site sequence composition of 25 ancestral nitrogenases (inferred across Tree-1–5) and all 284 extant nitrogenase sequences used for phylogenetic reconstruction. Our analysis shows clear distinctions between the active-site compositions of extant Mo-versus V-/Fe-nitrogenases (Figure 4a). Specifically, the mean identity between Vnf/Anf and Nif-I/Nif-II is ∼50%, as compared with the mean identity between Vnf and Anf (∼71%) and between Nif-I and Nif-II (∼77%). The difference between Vnf/Anf and Nif-I/Nif-II sequences is not seen as distinctly when comparing total HDK sequences (Figure 4b), suggesting that the active-site sequence differences, in particular, may be better indicators of metal dependence than whole sequence differences due to greater catalytic tuning toward the cofactor.

**Figure 4.**
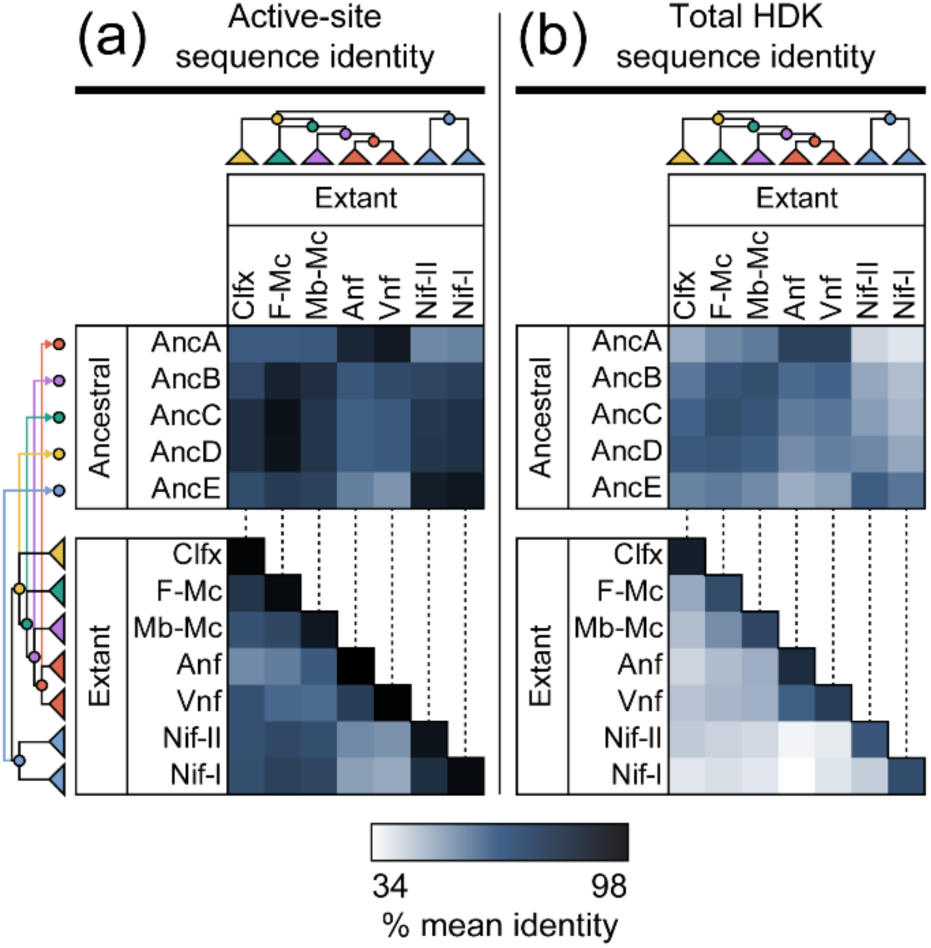
Active-site and full HDK sequence comparisons between extant and ancestral nitrogenases. (a) Active-site sequence identities of ancestral and extant nitrogenases. Active-site residues include 30 amino acids positioned within 5 Å of the active-site cofactor. (b) Total HDK sequence identities of ancestral and extant nitrogenases. Percentage identity values have been averaged within each field of comparison. All 25 maximum likelihood ancestors and 284 extant sequences included in this study were used for sequence identity calculation. Phylogeny coloring is as follows: Clfx, yellow; F-Mc, green; Mb-Mc, purple; Anf/Vnf, red; Nif-I/-II, blue.

Because we observed active-site sequence distinctions between extant Vnf/Anf and Nif-I/Nif-II nitrogenases, we compared active-site sequences of ancestral versus extant nitrogenases to provide clues regarding ancestral metal dependence. Nearly all ancestral nitrogenases, including those inferred for the oldest ancestral nodes, share greater active-site identity with Mo-nitrogenases than with V-/Fe-only-nitrogenases (Figure 4a). Mean active-site sequence identities between AncB– AncE and Anf/Vnf nitrogenases range between ∼50 and 63%, whereas those between AncB–AncE and Nif-I/Nif-II nitrogenases range between ∼69 and 87%. An exception is AncA (ancestral to Vnf and Anf), which has higher mean identity to Anf/Vnf nitrogenases (∼85%) than to Nif-I/Nif-II nitrogenases (∼51%). Because active-site sequence identity can reliably differentiate extant Mo-from V-/Fe-nitrogenases, the resemblance of most ancestral active sites to those of Mo-nitrogenases is suggestive of Mo-dependence. These results are consistent with previous sequence-based analyses that found that basal F-Mc nitrogenase sequences most resemble those of Mo-nitrogenases (McGlynn et al., 2012).

Finally, in order to incorporate the posterior probability distributions of inferred ancestral nitrogenase sequences into our sequence-based interpretations, we built a probabilistic model from the active-site amino acid probability distributions of 268 modern nitrogenases of known metal dependence (i.e., excepting 16 Clfx, F-Mc, and Mb-Mc homologs). Amino acid probability distributions were determined for active-site residues and used to train a support-vector classifier (see Section 2.2 for details). Surprisingly, the classifier was able to achieve 100% accurate classification on out-of-sample test data. However, principle-component analysis of active-site residue composition suggests that Nif sequences are completely separable from Vnf and Anf sequences using only one principle component, and Vnf and Anf sequences also separable using only two dimensions (Figure S6-1). The trained classifier was used to classify uncharacterized Clfx, F-Mc, and Mb-Mc sequences, as well as 25 maximum-likelihood ancestral nitrogenase sequences. All uncharacterized and ancestral sequences were classified as Nif-like, with the exception of all AncA ancestors, which were classified as Vnf-like (Table S6-1). Classification support (reported as distance of each data sample from the support-vector hyperplane defining the inferred class) for ancestral sequences is nearly identical across different ancestors inferred for the same phylogenetic node (Table S6-1). There is little difference in classification support between ancestors inferred from a MUSCLE alignment as from a MSAProbs alignment, with the exception between AncA-4 (inferred from MUSCLE alignment) and other AncA sequences (inferred from MSAProbs alignment). Support for AncA-4 classification is −0.0884 (for Nif), 2.1059 (for Vnf), and 0.9685 (for Anf), as opposed to mean support for other AncA sequences, 0.9774 ± 0.0023 (for Nif), 2.0626 ± 0.0056 (for Vnf), and −0.0455 ± 0.0046 (for Anf) (greater distance indicates greater support for the inferred classification). These probability-distribution-based classification results are consistent with the sequence-identity based comparisons, the latter of which similarly suggest higher identity between most uncharacterized/ancestral (AncB-E) sequences and extant Mo-nitrogenases than V-/Fe-nitrogenases. It is perhaps not surprising that the inclusion of the inferred ancestral probability distributions does not substantially change classifications of metal dependence, given that the support for active-site residues is high (>0.92), and that, of these, only five sites are reconstructed with plausible alternate states (alternate residues with probability >0.30) (See Section 3.3 for further details).

### 3.5 Active-site structural features are uninformative for inferring ancestral metal dependence

To investigate metal-specific features of ancestral nitrogenase structures, we generated homology models of both extant and ancestral nitrogenase D-subunits that house the active site (Figure 5a). First, we modeled 33 broadly sampled extant nitrogenase NifD, VnfD, and AnfD sequences to benchmark classifications of ancestral nitrogenase models. Second, we calculated structural models of 25 nitrogenase ancestors inferred by maximum likelihood and of 2,500 ancestors inferred by random Bayesian sampling of maximum likelihood site posterior probabilities (100 Bayesian samples per maximum likelihood ancestor). We generated ten model replicates per extant sequence and maximum likelihood sequence, and one model per Bayesian-sampled sequence. All structures were modeled with the FeMo-cofactor included (additional modeling runs were executed with the FeV-cofactor included; Appendix S3). In total, 3,080 models were generated with the FeMo-cofactor.

**Figure 5.**
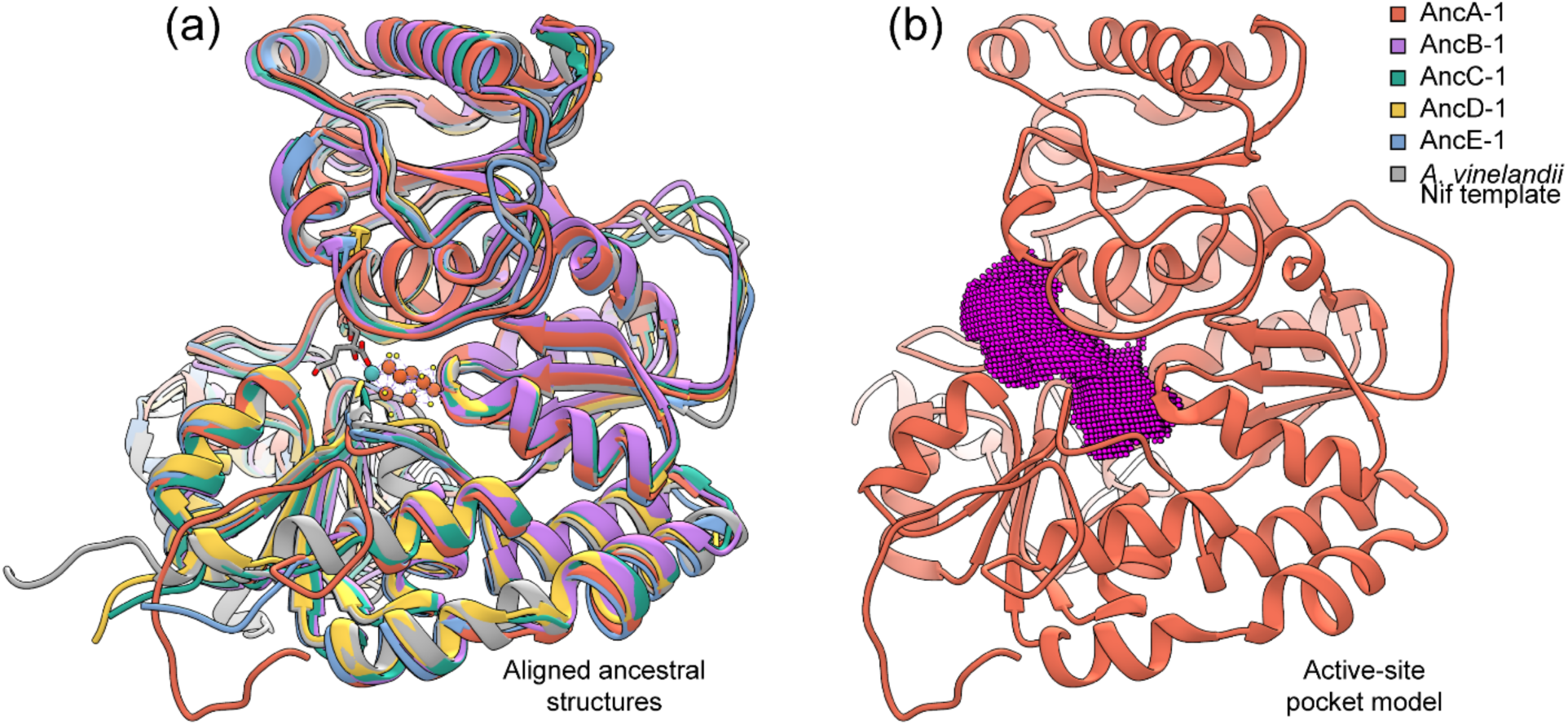
Structural and active-site pocket modelling of ancestral nitrogenases (a) Modeled D-subunit protein structures of ancestral nitrogenases inferred from the highest-likelihood phylogeny (Tree-1; Figure 2) aligned to an *A. vinelandii* Nif structural template (PDB 3U7Q (Spatzal et al., 2011)). (b) Example of a modeled active-site pocket for ancestral nitrogenase AncA-1. The 0.5-Å-resolution point field generated for pocket volume calculation is shown in pink.

For each of the 3,080 extant and ancestral D-subunit nitrogenase models, we calculated the volume of the active-site pocket (Figure 5b), a parameter previously used, in part, to classify the metal dependence of extant uncharacterized nitrogenases (McGlynn et al., 2012). These pocket volume values are plotted in Figure 6. Among modeled extant nitrogenases, mean pocket volumes are 1175.12 ± 51.93 Å^3^ for Mo-nitrogenases, 1121.86 ± 36.36 Å^3^ for V-nitrogenases, and 963.39 ± 75.80 Å^3^ for Fe-nitrogenases.

**Figure 6.**
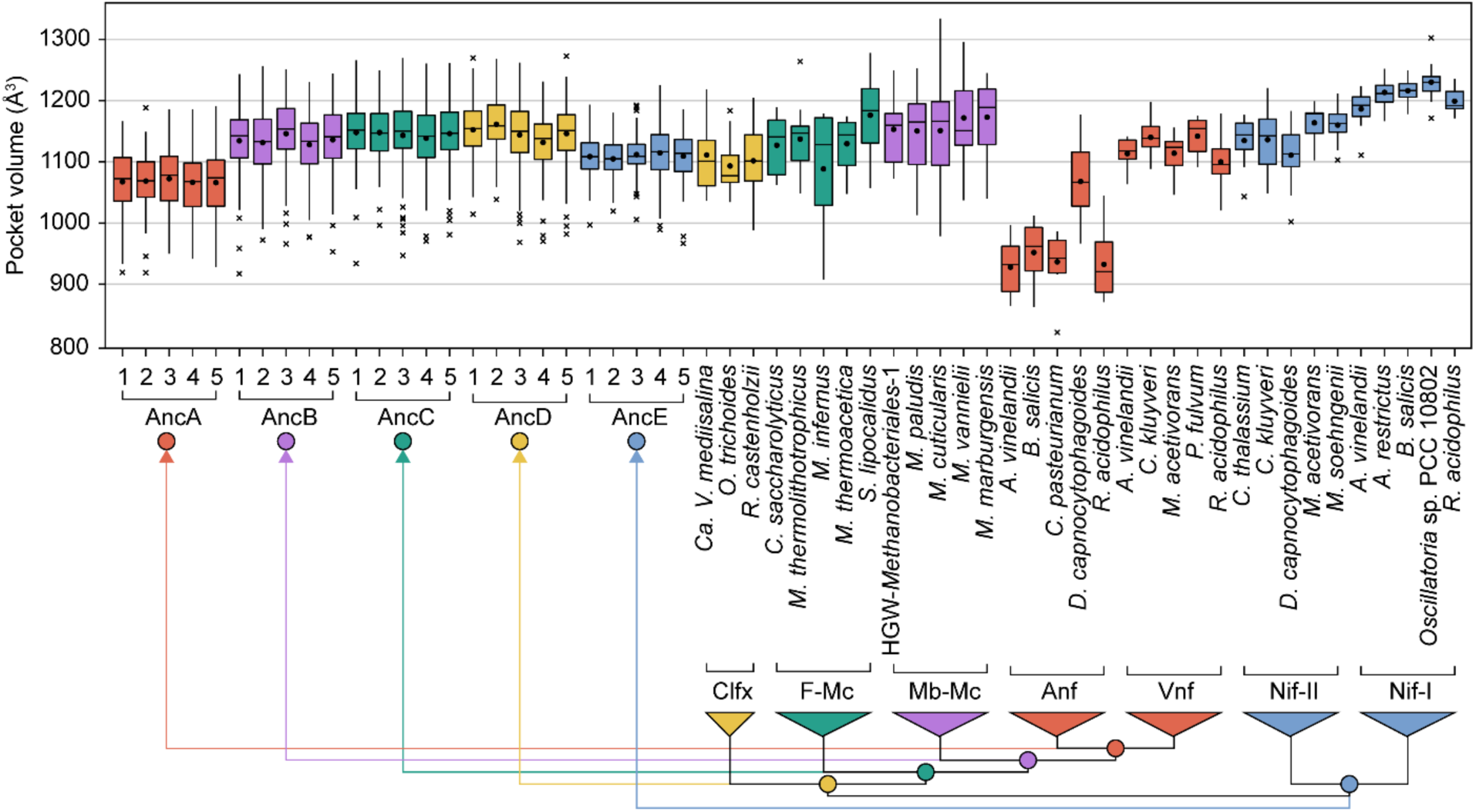
Extant and ancestral nitrogenase active-site pocket volumes. Pocket volumes calculated for ancestral and representative extant nitrogenase D-subunit structures modeled with the FeMo-cofactor. Each ancestral plot contains 110 volume calculations (ten model replicates per maximum likelihood sequence plus one model for each of 100 Bayesian-sampled sequences) and each extant plot contains 10 volume calculations (10 model replicates per extant sequence). Median values are indicated by bars, mean values by points, the range (excluding outliers) by whiskers, and outliers by crosses. Phylogeny coloring is as follows: Clfx, yellow; F-Mc, green; Mb-Mc, purple; Anf/Vnf, red; Nif-I/-II, blue.

We observe less difference between mean pocket volumes of extant V-nitrogenases (1121.86 ± 36.36 Å^3^) and Nif-II Mo-nitrogenases (1141.13 ± 46.30 Å^3^) than between Nif-I (1209.11 ± 30.79 Å^3^) and Nif-II Mo-nitrogenases. A statistical nonparametric test of volume median differences also suggests greater similarity between V- and Nif-II Mo-nitrogenases than between Nif-I and Nif-II nitrogenases (Appendix S3). All V-nitrogenase and Nif-II Mo-nitrogenase volume values range between 1002.13 and 1220.63 Å^3^, which problematically overlap with the volume ranges of AncB (917.38–1256.25 Å^3^), AncC (933.75–1269.50 Å^3^), AncD (968.50–1272.75 Å^3^), and AncE (966.75–1225.25 Å^3^) ancestral models, as well as ranges for uncharacterized Clfx (988.25– 1218.13 Å^3^) and F-Mc (908.00–1277.75 Å^3^) structures. The volume range of AncA models (919.00-1191.13 Å^3^) lies between the ranges of both V-(1020.75–1197.63 Å^3^) and Fe-nitrogenases (821.38–1177.00 Å^3^). These same overall patterns are observed when comparing only maximum likelihood ancestral models or Bayesian-sampled ancestral models, as well as alternate modeling runs with the FeV-cofactor (Appendix S3). Greater similarities between V- and Nif-II Mo-nitrogenases than between Nif-I and Nif-II Mo-nitrogenases suggests that, contrary to findings from previous analyses (McGlynn et al., 2012), modeled pocket volume is not informative for classifying modern or ancestral nitrogenases of unknown metal dependence. At the very least, the overlap in volume ranges between ancestors and extant isozymes of varying metal dependence in our analyses preclude the unambiguous classification of ancestral metal dependence by these structural features.

## 4 DISCUSSION

Nitrogenase mediates the reduction of N_2_ to NH_3_, a key step in nitrogen fixation (Hoffman et al., 2014; Seefeldt, Hoffman, & Dean, 2009). The metal dependence of nitrogenase, which impacts both catalytic properties (Eady, 1996; Harris, Yang, et al., 2018; Hu et al., 2011; Lee et al., 2018; Rebelein et al., 2018; Zheng et al., 2018) and ecological distribution (McRose, Zhang, Kraepiel, & Morel, 2017; Zhang et al., 2016), suggests a potential role for geochemical constraints on its evolution (Anbar & Knoll, 2002; Boyd, Hamilton, et al., 2011; Canfield et al., 2010; Raymond et al., 2004). Thus, understanding ancestral nitrogenase metal dependence can help resolve the early history of biological nitrogen fixation, and, in a broader sense, the impact that ancient metal availabilities have had on the evolution of biologically essential metabolisms over Earth history (Anbar & Knoll, 2002; Moore, Jelen, Giovannelli, Raanan, & Falkowski, 2017). Previous phylogenetic work has established that Mo-, V-, and Fe-nitrogenases, though genetically distinct, are homologous (Boyd, Anbar, et al., 2011; Boyd, Hamilton, et al., 2011; Raymond et al., 2004). Most recent phylogenetic analyses also indicate that V- and Fe-nitrogenases are derived from Mo-nitrogenases, the latter having originated subsequent to the gene duplication event that produced *nifE* and *nifN* (Boyd, Anbar, et al., 2011). However, the precise trajectory of metal-binding evolution in the nitrogenase family is not completely known, and discrepancies between current phylogenetics-based models and the geochemical record of nitrogen fixation (e.g., (Stueken et al., 2015)) remain.

We used phylogenetic reconstruction, ancestral sequence inference, and structural homology modeling to explore outstanding questions of early nitrogenase evolution and metal dependence. Though the tree reconstructions presented here are largely in congruence with previous phylogenetic analyses, certain topological differences, particularly with regard to basal uncharacterized homologs lacking one or more associated *nifEN* genes, suggest important deviations from previous narratives of early metal dependence (Boyd, Anbar, et al., 2011; Boyd, Hamilton, et al., 2011; Boyd & Peters, 2013; Raymond et al., 2004). The reconstruction of ancestral nitrogenase sequences *in silico* provides the means to directly infer ancient metal-binding properties from molecular information.

### 4.1 Inferred metal dependence of ancestral nitrogenases

The nitrogenase active site is known, in addition to the metal content of the cofactor, to contribute to the variable catalytic properties of different nitrogenase isozymes (Brigle et al., 1987; Christiansen et al., 2000; Fixen et al., 2016; Kim et al., 1995; Sarma et al., 2010; Yang et al., 2012). We therefore explored active-site features of ancestral nitrogenases for correlations with metal dependence.

We find that modeled active-site structural features are not informative for the inference of ancestral metal dependence. Pocket volumes of modeled extant nitrogenases do not appear to be strongly correlated with metal cofactor content. Problematically, the predicted volume ranges of oldest ancestors (as well as those of uncharacterized nitrogenases) overlap with both V- and Mo-nitrogenases (Figure 6). We acknowledge that these homology modeling results may not precisely reflect true biological differences (for which a comprehensive analysis is not yet possible due the limited availability of V- and Fe-nitrogenase structures). This is illustrated by the difference between pocket volumes of published structures and those of homology models (e.g., *A. vinelandii* PDB 3U7Q NifD structure pocket volume ≈ 994.750 and mean *A. vinelandii* NifD modeled structure pocket volume ≈ 1186.412). However, it is not surprising that active-site pocket volume does not predict metal dependence, given the probable similar sizes and structures of the FeMo-, FeV-, and FeFe-cofactors (Eady, 1996; Krahn et al., 2002; Daniel Sippel et al., 2018; Spatzal et al., 2011). These findings contrast with a previous study by McGlynn and coworkers (2012) in which pocket volume comparisons were used, in part, to classify the Mo-dependence of uncharacterized nitrogenases. In this previous study, the volume means for Mo-nitrogenases were sufficiently distinct from V- and Fe-nitrogenases as to provide unambiguous classification of Mo-dependence. Our analyses differ from this previous study in several ways: (1) we incorporated V-nitrogenase structural templates (D. Sippel & Einsle, 2017; Daniel Sippel et al., 2018) for homology modeling, not available at the time of the previous study, (2) we modeled 20 extant sequences of known metal dependence rather than 12, (3) we used consistent, explicit parameters for both homology modeling and pocket volume calculation across all sequences, (4) we included modeling replicates for pocket volume calculation. In summary, our expanded modeling analyses reveals a reduced efficacy of the pocket volume parameter for inference of ancestral (and uncharacterized) nitrogenase metal dependence.

Unlike structure-based analyses, we find that active-site sequence features do reliably differentiate Mo-nitrogenases and V-/Fe-nitrogenases, as was also previously noted by McGlynn and coauthors (2012). We find that both sequence-based and probability-distribution-based analyses are thus useful for classifying nitrogenases of unknown metal dependence, and are robust to the underlying uncertainty of ancestral sequence inference (Section 3.4; Appendix S6). Regarding AncA (ancestral to V- and Fe-nitrogenases), we identified specific active-site residues that have previously been suggested to interact with a proposed carbonate ligand not present in the FeMo-cofactor. These residues form a ^355^TGGPRL^360^ loop conserved only among V-nitrogenases and homologous to ^355^IGGLRP^360^ in *A. vinelandii* NifD (D. Sippel & Einsle, 2017). This substitution of Thr-355 for Ile-355, as well as the exchange of Leu and Pro positions may permit the inclusion of the FeV-cofactor carbonate ligand by VnfD that is not possible by the NifD protein (D. Sippel & Einsle, 2017). Thr-355 and Pro-358 are unique to V-nitrogenases, and Leu-360 is unique to V- and Fe-nitrogenases. All AncA sequences conserve the ^355^TGGPRL^360^ residue loop capable of accommodating the FeV-cofactor. Furthermore, AncA sequences generally exhibit greater numbers of residues unique to V-nitrogenases than those unique to Fe-nitrogenases, the mean identity of AncA active-site sequences is highest for V-nitrogenases (Figure 4a), and the classifier analysis based on active-site amino acid probability distributions suggests that AncA sequences most resemble modern V-nitrogenases (Section 3.4; Appendix S6). Together, these observations suggest that AncA is specific toward the VFe-cofactor.

Ancestral and extant sequence comparisons indicate that the active-sites of oldest nitrogenase ancestors (AncB–AncE) resemble those of extant Mo-nitrogenases more than V- or Fe-nitrogenases (Figure 4a; Appendix S6). This observation is particularly significant given that these same patterns are not observed across the total HDK sequence (Figure 4b). Specifically, this discrepancy supports the notion that the nitrogenase active-site has adapted to the catalytic properties of each metal cofactor over its evolutionary history (Harris, Yang, et al., 2018), and that this adaptation has manifested in active-site sequence differences that stand apart from baseline phylogenetic distance. Though we are not able to identify specific residues or motifs that may functionally be attributed to metal dependence (as with AncA), the resemblance of the early ancestral nitrogenase active site to those of Mo-nitrogenases is highly suggestive of an ancient role for an active-site cluster resembling the FeMo-cofactor.

### 4.2 A proposed model for the evolution of nitrogenase metal dependence over geologic time

Despite the lack of information provided by structural analyses, the active-site sequence features of oldest ancestral sequences (i.e., AncB–E) support the inference that these early nitrogenase incorporated the FeMo-cofactor or similar cluster. However, the observation that ancestral AncC and AncD active sites in particular most resemble those of extant Mo-nitrogenases is at odds with the phylogenetic distribution of *nifE* and *nifN* genes, which suggest that early-branching uncharacterized Clfx and F-Mc taxa (for which AncC and AncD are ancestral) do not possess the full biosynthetic complement in the canonical FeMo-cofactor assembly pathway.

The placement of Clfx and F-Mc clades in our analyses differs from previous phylogenetic reconstructions. For example, the phylogenetic tree presented by Boyd and coauthors nests Clfx and F-Mc clades within Mo-nitrogenases, which notably branch more recently than V-, Fe-, and Mb-Mc Mo-nitrogenases (Boyd, Hamilton, et al., 2011). Such a topology would be consistent with the secondary loss of *nifEN* genes in Clfx and F-Mc clades, and the absence of such genes would not likely represent an ancestral state for the cofactor biosynthetic pathway. By contrast, we find that Clfx and F-Mc clades are early-branching, and that the presence of *nifE* and *nifN* genes decreases stepwise with divergence age within the uncharacterized/V-/Fe-nitrogenase lineage: Mb-Mc taxa, most recently branched, have both *nifEN,* most F-Mc taxa only have *nifE*, and Clfx taxa, earliest branched, have neither (*vnf* and *anf* gene clusters, resulting only from the duplication of *nif* structural genes (Boyd, Hamilton, et al., 2011), typically lack dedicated scaffolding genes and instead co-opt *nifEN;* (Hamilton et al., 2011; Mus et al., 2018)). One may thus parsimoniously conclude that uncharacterized AncC–D ancestors similarly lacked one or more *nifEN* genes. A more recent, previously published topology is more similar to the tree presented here, though lacking in Clfx sequences and thus missing relevant information regarding the lack of associated biosynthetic components for early-diverged homologs (Boyd & Peters, 2013). It is possible that the larger and more broadly sampled sequence dataset used here has refined the placement of these uncharacterized clades, which is supported by our analyses with an expanded outgroup that maintains the positions of Clfx and F-Mc sequences (Appendix S2).

At least two possible models might resolve the discrepancy between inferred early Mo-dependence and suggested lack of cofactor biosynthesis genes: (1) The last common nitrogenase ancestor represented in our tree was in fact hosted by an organism that possessed *nifEN* genes and was able to synthesize the FeMo-cofactor via the full, canonical biosynthetic pathway. Further, one or more *nifEN* genes were lost in both Clfx and F-Mc taxa, but were retained in all ancestors (AncA–E) as well as in Nif-I, Nif-II, and Mb-Mc taxa. The resemblance of oldest ancestral active sites (AncB–E) to those of Mo-nitrogenases indicate ancient FeMo-cofactor dependence, and the same resemblance of Clfx and F-Mc nitrogenases to Mo-nitrogenases may be inherited from these ancestors. (2) The last common nitrogenase ancestor was hosted by an organism that did not possess *nifEN* genes, and achieved cofactor biosynthesis by alternative and/or simplified means. Such an early-evolved and perhaps inefficient pathway for cofactor synthesis may still be present in F-Mc taxa, and possibly Clfx taxa as well. *nifEN* genes then evolved through the uncharacterized/V-/Fe-nitrogenase lineage via gene duplication of *nifDK* and subsequent subfunctionalization (Boyd, Anbar, et al., 2011), and was selected for once the canonical FeMo-cofactor pathway was established prior to the AncB nitrogenase at the earliest, but at least in Mb-Mc nitrogenases (*nifEN* genes may have been transferred between ancestors of Mb-Mc, Nif-I, and Nif-II taxa).

We prefer the second model for several reasons:

1. It is evolutionarily unlikely for a portion of the FeMo-cofactor biosynthetic pathway to be lost in Clfx and F-Mc taxa. To our knowledge, no other Mo-nitrogenases are known to lack associated *nifEN* genes, suggesting that the canonical biosynthetic pathway for the FeMo-cofactor is selectively advantageous (*anf* and *vnf* gene clusters typically lack dedicated scaffolding genes and co-opt *nifEN*; (Hamilton et al., 2011; Mus et al., 2018)). Rather, it is more parsimonious to infer that these early-branching uncharacterized clades diverged prior to the *nifDK* duplication that produced *nifEN*, as is evidenced by the nesting of *nifEN* genes within *nifDK* clades in previous phylogenetic reconstructions (Boyd, Anbar, et al., 2011).
2. It has previously been proposed that early nitrogenases may have been capable of reducing nitrogen prior to the development of the canonical FeMo-cofactor biosynthetic pathway (Boyd, Hamilton, et al., 2011; Boyd & Peters, 2013; Mus, Colman, Peters, & Boyd, 2019; Soboh, Boyd, Zhao, Peters, & Rubio, 2010). Further, an ancient Mo-independent nitrogenase may have been capable of — albeit inefficiently — reducing nitrogen by a cofactor resembling the Fe-S-C cluster assembled by NifB, which constitutes the biosynthetic precursor to the FeMo-cofactor (Boyd & Peters, 2013; Mus et al., 2019; Soboh et al., 2010). Though our sequence analyses cannot fully assess ancestral nitrogenase dependence for a NifB-cofactor, it is likely that the NifB-cofactor resembles the structure and composition of the FeFe-cofactor, excepting homocitrate (Corbett et al., 2006; Guo et al., 2016; Harris, Lukoyanov, et al., 2018). However, the greater similarity of AncC–D active sites to those of Mo-nitrogenases than Fe-nitrogenases likely suggests ancestral dependence on a cofactor incorporating Mo rather than only Fe, as would be the case for the NifB-cofactor. Instead, it is possible that, lacking *nifEN*, an alternative and/or simplified pathway for FeMo- or similar cofactor synthesis may have acted as a transition state between such an Mo-independent stage (i.e., binding a cluster resembling the NifB-cofactor; (Boyd & Peters, 2013; Mus et al., 2019)) and the development of the canonical FeMo-cofactor biosynthetic pathway. It is thus reasonable to speculate that this transition to Mo-usage may be exhibited by AncC–D ancestors.
3. Mo-nitrogenases are far more efficient at reducing nitrogen than other isozymes (Eady, 1996; Harris et al., 2019; Harris, Yang, et al., 2018), and the majority of all extant nitrogenases are Mo-dependent across both anoxic and oxic environments (Boyd et al., 2015; Mus et al., 2019; Raymond et al., 2004). Even those organisms that have additional V- or Fe-nitrogenases still retain and preferentially express Mo-nitrogenases (Boyd, Anbar, et al., 2011; Boyd, Hamilton, et al., 2011; Dos Santos et al., 2012; Hamilton et al., 2011; Raymond et al., 2004). The presence of a pathway, albeit not fully established, for FeMo-cofactor or similar cofactor synthesis would be selectively advantageous even if Mo was transiently available prior to ∼2.3–2.5 Ga Earth surface oxygenation (Anbar et al., 2007; Anbar & Knoll, 2002; Canfield et al., 2010; Lyons et al., 2014; Raymond et al., 2004). In fact, previous studies suggest an early generalist behavior for ancient nitrogenases with regard to their metal usage (Raymond et al., 2004). Such limited and possibly non-specific Mo-incorporation into an active-site cluster early in nitrogenase history could pave the way for increased Mo-selectivity over time via the addition of NifEN biosynthetic components, building from an earlier Mo-independent state incorporating something resembling the modern NifB-cofactor (Boyd & Peters, 2013; Mus et al., 2019). An early nitrogenase that was effective at incorporating Mo would have favored the selection of other ancillary genes that later gave rise to increased Mo-specificity.
4. ∼3.2-Ga nitrogen isotopic signatures have been interpreted to reflect ancient Mo-dependent nitrogenase activity, due to similarities in isotopic fractionation by extant Mo-nitrogenases (Stueken et al., 2015). However, this interpretation conflicts with age estimates of both *nifEN* (inferred to also reflect the age of the FeMo-cofactor) (Boyd, Anbar, et al., 2011) and of earliest marine Mo availability (Anbar et al., 2007; Lyons et al., 2014). It is possible that an alternative pathway for FeMo- of similar cofactor synthesis in early nitrogenases may provide an explanation for these so-far unresolved, early isotopic signatures.

The proposed model of an ancestral, alternative pathway for Mo-cofactor assembly, mapped to our phylogenetic reconstruction and inferred ancestors, is illustrated in Figure 7. In the first stage, represented by AncD and extant Clfx homologs, ancient enzymes use an alternative and/or simplified pathway to incorporate an Mo-containing cluster, likely resembling or possibly identical to the canonical FeMo-cofactor in the absence of NifEN. So as to remain agnostic toward the particular cofactor structure, we term this a “proto-Mo-cofactor” (Figure 7a). Such enzymes may not be capable of reducing nitrogen, as this has not been demonstrated definitively for Clfx homologs. In the absence of both NifEN, cofactor assembly may be inefficient and subject to transient marine Mo availabilities prior to ∼2.3 – 2.5 Ga Earth surface oxygenation, as evidenced by geochemical data (Anbar et al., 2007). In the second stage, represented by AncC and extant F-Mc nitrogenases, gene duplication of NifD forms NifE, which may be able to form a homodimer or homotetramer for the scaffolding of the FeMo-cofactor (Figure 7b). In the third stage, represented by AncB, AncE, and extant Mb-Mc nitrogenases, full gene duplication of NifDK additionally forms NifN (Figure 7c). Together with NifB, NifE and NifN complete the canonical FeMo-cofactor biosynthetic pathway, resulting in more refined nitrogenase Mo-dependence. In the fourth stage, gene duplication of *nif* results in, first, *vnf*, followed by *anf* (Figure 7d). These V- and Fe-dependent enzymes rely on NifBEN biosynthetic components (though some V-nitrogenases, as in *A. vinelandii*, rely on more recently evolved VnfEN proteins that derive from a duplication of NifEN; (Boyd, Anbar, et al., 2011)). Though the greater efficiency of Mo-nitrogenase results in widespread diversification, V- and Fe-nitrogenases provide selective advantage in microenvironments deficient in Mo. This model, built from the phylogenetic and ancestral sequence inferences provided here, as well as from decades of previous geobiological investigations of nitrogenase evolution, helps resolve outstanding questions regarding ancient metal dependence, and, importantly, provides testable hypotheses for future investigations. Such investigations may, for example, seek to clarify the nitrogen capability and metal dependence of uncharacterized Clfx and F-Mc nitrogenases, as well as experimentally resurrect and characterize ancestral nitrogenases in the laboratory.

**Figure 7.**
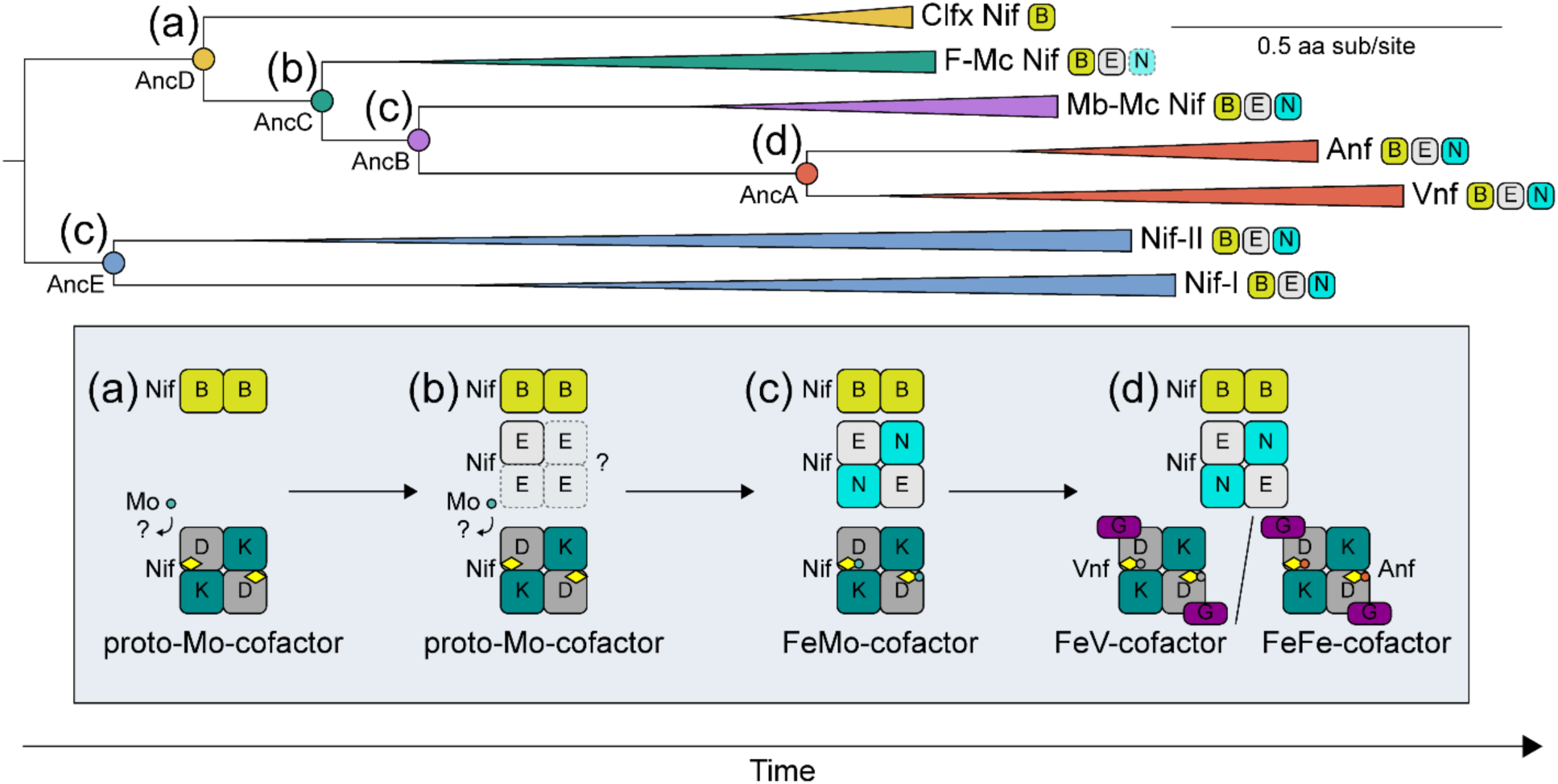
Proposed model for the evolution of nitrogenase metal specificity from an ancestral, alternative and/or simplified pathway for Mo-cofactor incorporation. Full description of stages (a)–(d) in the evolution of nitrogenase metal dependence is provided in the main text (Section 4.2). Possible alternative pathway for nitrogenase Mo-cofactor synthesis indicated in stages (a)–(b). Phase (a)–(d) are also mapped to analyzed ancestors within the nitrogenase phylogeny. Presence of NifB, NifE, and NifN in represented taxa indicated next to each clade of the phylogeny (F-Mc clade has only one species that harbors NifN).

## CONCLUSION

We reconstructed the phylogenetic history of nitrogenase proteins, as well as inferred and modelled ancestral nitrogenase sequences, in order to explore the evolutionary trajectory of nitrogenase metal dependence. We find that, whereas modeled structural features of ancestral nitrogenases do not offer conclusive indications of ancient metal usage, active-site sequence features of oldest ancestors most resemble those of extant Mo-nitrogenases. The absence of associated cofactor biosynthesis proteins that function within the canonical FeMo-cofactor biosynthetic pathway in several early-branching, uncharacterized homologs evidences a possible alternative and/or simplified pathway for Mo-cofactor assembly. We speculate that this alternative pathway may today be present in extant uncharacterized taxa, and we propose a model wherein canonical Mo-cofactor assembly evolved via the stepwise introduction of FeMo-cofactor biosynthetic components following the divergence of more basal uncharacterized lineages. V-nitrogenases subsequently diversified, followed by Fe-nitrogenases, in agreement with previous phylogenetic inferences that Mo-dependence evolved first (Boyd, Hamilton, et al., 2011). This model helps to reconcile phylogenetic and geobiological explanations of nitrogenase evolution (Anbar & Knoll, 2002; Boyd, Anbar, et al., 2011; Boyd, Hamilton, et al., 2011). Future molecular paleobiology studies, particularly those that integrate experimental assessments of laboratory-resurrected ancestral nitrogenases, may continue to refine our understanding of nitrogenase and environmental coevolution.

## Supporting information

Complete SI List

## ACKNOWLEDGEMENTS

This work was supported by a NASA Astrobiology Postdoctoral Fellowship (Garcia), the Harvard Origins of Life Initiative (Kacar, McShea), the National Science Foundation (Kacar, #1724090), the John Templeton Foundation (Kacar, #61239), a NASA Early Career Fellowship (Kacar, 80NSSC19K1617), and the University of Arizona Foundation Small Grants Program (Kacar, Garcia). We thank the Kacar Lab, Scott Edwards, Jennifer Glass, Andrew Knoll, Sebastian Kopf, Lance Seefeldt, and Xinning Zhang for invaluable discussions, as well as three anonymous reviewers for constructive feedback.

