## Supplemental File 1 for "Reconstructing the Evolutionary History of Nitrogenases: Evidence for Ancestral Molybdenum-Cofactor Utilization"

**Table S1.** List of extant nitrogenase and outgroup protein sequences used for phylogenetic reconstruction and ancestral sequence inference.

| **Species** | **Accession/Locus Tag** | | | **Phylum** | **Class** |
| --- | --- | --- | --- | --- | --- |
|  | **NifH** | **NifD** | **NifK** |  |  |
| Acetobacterium woodii | WP_014355191 | WP_014355188 | WP_014355187 | Firmicutes | Clostridia |
| Acidiferrobacter thiooxydans | WP_065969940 | WP_065969942 | WP_065969944 | Proteobacteria | Gammaproteobacteria |
| Acidihalobacter prosperus | WP_070071968 | WP_070073945 | WP_070071969 | Proteobacteria | Gammaproteobacteria |
| Acidithiobacillus ferrooxidans | WP_113525799 | WP_113525798 | WP_113525797 | Proteobacteria | Acidithiobacillia |
| Actinobacteria bacterium HGW-Actinobacteria-10 | PKQ29728 | PKQ29729 | PKQ29730 | Actinobacteria | - |
| Afifella marina | WP_092811147 | WP_092811149 | WP_092811151 | Proteobacteria | Alphaproteobacteria |
| Agarivorans albus | WP_016403298 | WP_016403297 | WP_016403296 | Proteobacteria | Gammaproteobacteria |
| Alkalibacter saccharofermentans | WP_073269025 | WP_073269336 | WP_073269028 | Firmicutes | Clostridia |
| Anaerobacillus alkalilacustris | WP_071308028 | WP_071308027 | WP_071308026 | Firmicutes | Bacilli |
| Anaeromyxobacter sp. K | WP_012527198 | WP_012527199 | WP_012527200 | Proteobacteria | Deltaproteobacteria |
| Anaerovirgula multivorans | WP_089285040 | WP_089282036 | WP_089282035 | Firmicutes | Clostridia |
| Aneurinibacillus terranovensis | WP_027417121 | WP_027417120 | WP_027417119 | Firmicutes | Bacilli |
| Arcticibacter svalbardensis | WP_016195754 | WP_016195751 | WP_040299905 | Bacteroidetes | Sphingobacteriia |
| Azohydromonas australica | WP_029001236 | WP_029001237 | WP_029001238 | Proteobacteria | Betaproteobacteria |
| Azomonas agilis DSM 375 | LX59DRAFT_00350 | LX59DRAFT_00351 | LX59DRAFT_00352 | Proteobacteria | Gammaproteobacteria |
| Azospira sp. I13 | WP_109041643 | WP_109041644 | WP_109041645 | Proteobacteria | Betaproteobacteria |
| Azospirillum halopraeferens | WP_029011025 | WP_029011024 | WP_029011023 | Proteobacteria | Alphaproteobacteria |
| Azotobacter vinelandii DJ, ATCC BAA-1303 | Avin_01380 | Avin_01390 | Avin_01400 | Proteobacteria | Gammaproteobacteria |
| Azovibrio restrictus | WP_026687661 | WP_026687660 | WP_026687659 | Proteobacteria | Betaproteobacteria |
| Bacillus caseinilyticus | WP_090885364 | WP_090885366 | WP_090885368 | Firmicutes | Bacilli |
| Beijerinckia indica | WP_012383484 | WP_012383485 | WP_012383486 | Proteobacteria | Alphaproteobacteria |
| Betaproteobacteria bacterium HGW-Betaproteobacteria-11 | PKO84828 | PKO84829 | PKO84997 | Proteobacteria | Betaproteobacteria |
| Blastopirellula marina | WP_105335631 | WP_105335632 | WP_105335633 | Planctomycetes | Planctomycetia |
| Bradyrhizobium diazoefficiens USDA 110 | blr1769 | blr1743 | blr1744 | Proteobacteria | Alphaproteobacteria |
| Brenneria salicis ATCC 15712 | Bresa_01966 | Bresa_01965 | Bresa_01964 | Proteobacteria | Gammaproteobacteria |
| Caenispirillum bisanense | WP_097278677 | WP_097278676 | WP_097278675 | Proteobacteria | Alphaproteobacteria |
| Caldicellulosiruptor saccharolyticus | WP_011917964 | WP_011917961 | WP_011917960 | Firmicutes | Clostridia |
| Calditerrivibrio nitroreducens | WP_013450808 | WP_013450809 | WP_013450810 | Deferribacteres | Deferribacteres |
| Candidatus Atelocyanobacterium thalassa isolate ALOHA | WP_012954002 | WP_012954003 | WP_012954004 | Cyanobacteria | - |
| Candidatus Lambdaproteobacteria bacterium RIFOXYC1_FULL_56_13 | OGH03497 | OGH03496 | OGH03495 | Proteobacteria | Candidatus Lambdaproteobacteria |
| Candidatus Lambdaproteobacteria bacterium RIFOXYD2_FULL_50_16 | OGG94501 | OGG94500 | OGG94499 | Proteobacteria | Candidatus Lambdaproteobacteria |
| Candidatus Magnetoovum chiemensis | KJR40986 | KJR40987 | KJR40988 | Nitrospirae | Nitrospira |
| Candidatus Margulisbacteria bacterium GWD2_39_127 | OGH94801 | OGH94911 | OGH94798 | Candidatus Margulisbacteria | - |
| Candidatus Marispirochaeta associata | WP_069897282 | WP_069897284 | WP_069897285 | Spirochaetes | Spirochaetia |
| Candidatus Thiodiazotropha endolucinida | WP_069124654 | WP_069124653 | WP_069124652 | Proteobacteria | Gammaproteobacteria |
| Candidatus Viridilinea mediisalina | WP_097642292 | WP_097642294 | WP_097642295 | Chloroflexi | Chloroflexia |
| Carboxydocella thermautotrophica | WP_078665332 | WP_078665331 | WP_078665330 | Firmicutes | Clostridia |
| Cellulosilyticum lentocellum | WP_013655494 | WP_013655497 | WP_013655498 | Firmicutes | Clostridia |
| Cereibacter changlensis | WP_107662511 | WP_107662510 | WP_107662509 | Proteobacteria | Alphaproteobacteria |
| Chlorobium limicola | WP_012465649 | WP_012465646 | WP_012465645 | Chlorobi | Chlorobia |
| Chloroherpeton thalassium ATCC 35110 | Ctha_1035 | Ctha_1032 | Ctha_1031 | Chlorobi | Chlorobia |
| Chroococcidiopsis sp. TS-821 | WP_104545041 | WP_104545042 | WP_104545043 | Cyanobacteria | - |
| Chrysiogenes arsenatis | WP_027390771 | WP_027390770 | WP_027390769 | Chrysiogenetes | Chrysiogenetes |
| Clostridiales bacterium DRI-13 | WP_034423366 | WP_034423373 | WP_034423375 | Firmicutes | Clostridia |
| Clostridium kluyveri DSM 555 | CKL_3081 | CKL_3078 | CKL_3077 | Firmicutes | Clostridia |
| Cohaesibacter sp. ES.047 | WP_096174773 | WP_096174774 | WP_096174775 | Proteobacteria | Alphaproteobacteria |
| Consotaella salsifontis | WP_078709045 | WP_078709046 | WP_078709047 | Proteobacteria | Alphaproteobacteria |
| Coraliomargarita akajimensis | WP_013044574 | WP_013044573 | WP_013044572 | Verrucomicrobia | Opitutae |
| Cupriavidus sp. amp6 | WP_029049998 | WP_029049999 | WP_029050000 | Proteobacteria | Betaproteobacteria |
| Cylindrospermopsis raciborskii | WP_061544850 | WP_061544851 | WP_061544852 | Cyanobacteria | - |
| Dechloromonas aromatica | WP_011287173 | WP_011287172 | WP_011287171 | Proteobacteria | Betaproteobacteria |
| Defluviitalea phaphyphila | WP_058485536 | WP_058485595 | WP_058485533 | Firmicutes | Clostridia |
| Dehalococcoides mccartyi | WP_010936850 | WP_010936847 | WP_010936846 | Chloroflexi | Dehalococcoidia |
| Dehalogenimonas sp. WBC-2 | AKG53288 | AKG53285 | AKG53284 | Chloroflexi | Dehalococcoidia |
| Deltaproteobacteria bacterium HGW-Deltaproteobacteria-4 | PKN11897 | PKN11885 | PKN11884 | Proteobacteria | Deltaproteobacteria |
| Dendrosporobacter quercicolus | WP_092073275 | WP_092073284 | WP_092073287 | Firmicutes | Negativicutes |
| Denitrovibrio acetiphilus | WP_013010353 | WP_013010354 | WP_013010355 | Deferribacteres | Deferribacterales |
| Derxia gummosa | WP_028310376 | WP_028310375 | WP_028310374 | Proteobacteria | Betaproteobacteria |
| Desertifilum sp. IPPAS B-1220 | WP_069969273 | WP_069969274 | WP_069969275 | Cyanobacteria |  |
| Desulfarculus baarsii 2st14, DSM 2075 | Deba_0443 | Deba_0440 | Deba_0439 | Proteobacteria | Deltaproteobacteria |
| Desulfatibacillum alkenivorans AK-01 | Dalk_1522 | Dalk_1519 | Dalk_1518 | Proteobacteria | Deltaproteobacteria |
| Desulfitobacterium hafniense Y51 | WP_005813529 | WP_005813531 | WP_011461720 | Firmicutes | Clostridia |
| Desulfobacca acetoxidans ASRB2, DSM 11109 | Desac_0350 | Desac_0353 | Desac_0354 | Proteobacteria | Deltaproteobacteria |
| Desulfobacterium autotrophicum HRM2, DSM 3382 | HRM2_09670 | HRM2_09700 | HRM2_09710 | Proteobacteria | Deltaproteobacteria |
| Desulfonatronum lacustre | WP_028571559 | WP_028571556 | WP_028571555 | Proteobacteria | Deltaproteobacteria |
| Desulforegula conservatrix | WP_027358827 | WP_027358824 | WP_027358823 | Proteobacteria | Deltaproteobacteria |
| Desulfosarcina cetonica | WP_054697558 | WP_054697549 | WP_054697545 | Proteobacteria | Deltaproteobacteria |
| Desulfosporosinus sp. OT | DOT_2734 | DOT_2733 | DOT_2732 | Firmicutes | Clostridia |
| Desulfotomaculum copahuensis | WP_066665790 | WP_066665786 | WP_066665785 | Firmicutes | Clostridia |
| Desulfovibrio sp. 86-1 | Ga0302701_112362 | Ga0302701_112359 | Ga0302701_112358 | Proteobacteria | Deltaproteobacteria |
| Desulfuribacillus alkaliarsenatis | WP_069643779 | WP_069643776 | WP_069643775 | Firmicutes | Bacilli |
| Desulfurispora thermophila | WP_018085363 | WP_018085366 | WP_018085367 | Firmicutes | Clostridia |
| Desulfurivibrio alkaliphilus AHT2 | DaAHT2_2455 | DaAHT2_2458 | DaAHT2_2459 | Proteobacteria | Deltaproteobacteria |
| Desulfurobacterium atlanticum | WP_089322897 | WP_089322840 | WP_089322896 | Aquificae | Desulfurobacteriales |
| Desulfuromonas soudanensis | WP_053552385 | WP_053551332 | WP_053551331 | Proteobacteria | Deltaproteobacteria |
| Desulfuromusa kysingii | WP_092350255 | WP_092350253 | WP_092350324 | Proteobacteria | Deltaproteobacteria |
| Dethiobacter alkaliphilus | WP_008518850 | WP_008518845 | WP_008518843 | Firmicutes | Clostridia |
| Dethiosulfatibacter aminovorans | WP_073047220 | WP_073047054 | WP_073047056 | Firmicutes | Tissierellia |
| Dickeya dadantii Ech703 | Dd703_0491 | Dd703_0492 | Dd703_0493 | Proteobacteria | Gammaproteobacteria |
| Dissulfuribacter thermophilus | WP_067619323 | WP_067619329 | WP_067619330 | Proteobacteria | Deltaproteobacteria |
| Draconibacterium sediminis | WP_045033459 | WP_045033456 | WP_045033455 | Bacteroidetes | Bacteroidia |
| Dysgonomonas capnocytophagoides | WP_026626156 | WP_051290673 | WP_026626160 | Bacteroidetes | Bacteroidia |
| Ectothiorhodospira marina | WP_090250046 | WP_090250049 | WP_090250051 | Proteobacteria | Gammaproteobacteria |
| Elusimicrobia RIFOXYB2_FULL_49_7 | OGS33428 | OGS33426 | OGS33425 | Elusimicrobia | - |
| Ethanoligenens harbinense YUAN-3T, DSM 18485 | Ethha_1567 | Ethha_1564 | Ethha_1563 | Firmicutes | Clostridia |
| Faecalicatena contorta | WP_109714170 | WP_109714176 | WP_109714178 | Firmicutes | Clostridia |
| Firmicutes bacterium HGW-Firmicutes-14 | PKM81963 | PKM81966 | PKM81967 | Firmicutes | - |
| Firmicutes bacterium HGW-Firmicutes-2 | PKM68346 | PKM68349 | PKM68350 | Firmicutes | - |
| Fischerella sp. PCC 9605 | WP_026731053 | WP_026731054 | WP_026731055 | Cyanobacteria | - |
| Fontibacillus panacisegetis DSM 28129 | Ga0074847_109103 | Ga0074847_109102 | Ga0074847_109101 | Firmicutes | Bacilli |
| Frankia casuarinae | WP_063578012 | WP_063577989 | WP_063577990 | Actinobacteria | Actinobacteria |
| Gallionellales GWA2_59_43 | OGS92466 | OGS92467 | OGS92468 | Proteobacteria | Betaproteobacteria |
| Gallionellales GWA2_60_18 | OGS90295 | OGS90408 | OGS90294 | Proteobacteria | Betaproteobacteria |
| Gammaproteobacteria bacterium 28-57-27 | OYY73708 | OYY73707 | OYY73706 | Proteobacteria | Gammaproteobacteria |
| Gammaproteobacteria bacterium HGW-Gammaproteobacteria-1 | PKM46060 | PKM46061 | PKM46062 | Proteobacteria | Gammaproteobacteria |
| Gammaproteobacteria bacterium RIFOXYD12_FULL_61_37 | OGT91538 | OGT91539 | OGT91540 | Proteobacteria | Gammaproteobacteria |
| Geminisphaera colitermitum | WP_043581909 | WP_081721723 | WP_043581908 | Verrucomicrobia | Opitutae |
| Geobacter metallireducens | WP_004514270 | WP_004514271 | WP_004514272 | Proteobacteria | Deltaproteobacteria |
| Geobacteraceae bacterium GWC2_53_11 | OGU17931 | OGU17930 | OGU17929 | Proteobacteria | Deltaproteobacteria |
| Geofilum rubicundum | WP_062122350 | WP_062122347 | WP_062122346 | Bacteroidetes | Bacteroidia |
| Geopsychrobacter electrodiphilus | WP_020677891 | WP_020677892 | WP_020677893 | Proteobacteria | Deltaproteobacteria |
| Geothermobacter sp. EPR-M | WP_085011015 | WP_085010978 | WP_085010979 | Proteobacteria | Deltaproteobacteria |
| Geovibrio sp. L21-Ace-BES | WP_022850366 | WP_022850365 | WP_022850364 | Deferribacteres | Deferribacterales |
| Gorillibacterium timonense SN4 | Ga0112892_17330 | Ga0112892_17329 | Ga0112892_17328 | Firmicutes | Bacilli |
| Gracilibacter sp. BRH_c7a | KUO65809 | KUO65806 | KUO65805 | Firmicutes | Clostridia |
| Halanaerobium saccharolyticum | WP_108140022 | WP_108139986 | WP_108139984 | Firmicutes | Clostridia |
| Halothece sp. PCC 7418 | WP_015225829 | WP_015225830 | WP_015225831 | Cyanobacteria | - |
| Halothiobacillus sp. LS2 | WP_066098062 | WP_066098061 | WP_066098060 | Proteobacteria | Gammaproteobacteria |
| Hartmannibacter diazotrophicus | WP_099558078 | WP_099558077 | WP_099558076 | Proteobacteria | Alphaproteobacteria |
| Heliobacterium modesticaldum | WP_012282218 | WP_012282219 | WP_012282220 | Firmicutes | Clostridia |
| Herbaspirillum seropedicae SmR1 | Hsero_2853 | Hsero_2852 | Hsero_2851 | Proteobacteria | Betaproteobacteria |
| Holophaga foetida | WP_005034263 | WP_005034268 | WP_005034270 | Acidobacteria | Holophagae |
| Hungateiclostridium cellulolyticum | WP_010251784 | WP_010251793 | WP_010251796 | Firmicutes | Clostridia |
| Hydrococcus rivularis | WP_073598931 | WP_073599038 | WP_073598932 | Cyanobacteria | - |
| Hydrogenophaga flava | WP_066257627 | WP_066257625 | WP_066257623 | Proteobacteria | Betaproteobacteria |
| Hydrogenophilales bacterium 28-61-23 | OYY93902 | OYY93903 | OYY93904 | Proteobacteria | Hydrogenophilalia |
| Ilyobacter polytropus CuHBu1, DSM 2926 | Ilyop_1410 | Ilyop_1407 | Ilyop_1406 | Fusobacteria | Fusobacteriia |
| Immundisolibacter cernigliae | WP_068802292 | WP_068802293 | WP_068802294 | Proteobacteria | Gammaproteobacteria |
| Labilibaculum manganireducens | WP_101310083 | WP_101310080 | WP_101310079 | Bacteroidetes | Bacteroidia |
| Lamprocystis purpurea | WP_020505238 | WP_020505237 | WP_020505236 | Proteobacteria | Gammaproteobacteria |
| Lebetimonas sp. JS032 | WP_024791553 | WP_024791554 | WP_024791555 | Proteobacteria | Epsilonproteobacteria |
| Lentisphaerae bacterium GWF2_50_93 | OGV59092 | OGV59114 | OGV59095 | Lentisphaerae | - |
| Lentisphaerae bacterium GWF2_57_35 | OGV45350 | OGV45347 | OGV45346 | Lentisphaerae | - |
| Leptolyngbya ohadii | WP_088891457 | WP_088891456 | WP_088891455 | Cyanobacteria | - |
| Leptospirillum ferriphilum | WP_099590653 | WP_099590654 | WP_099590466 | Nitrospirae | Nitrospira |
| Leptothrix cholodnii SP-6 | Lcho_1430 | Lcho_1431 | Lcho_1432 | Proteobacteria | Betaproteobacteria |
| Lutibacter agarilyticus | WP_089382364 | WP_089382367 | WP_089382368 | Bacteroidetes | Flavobacteriia |
| Lyngbya aestuarii | WP_023065299 | WP_023065312 | WP_023065341 | Cyanobacteria | - |
| Magnetococcus marinus MC-1 | Mmc1_1202 | Mmc1_1201 | Mmc1_1200 | Proteobacteria | Alphaproteobacteria |
| Magnetofaba australis | WP_085442124 | WP_085442125 | WP_085442126 | Proteobacteria | Alphaproteobacteria |
| Magnetospirillum bellicus VDY | MagVDY_03872 | MagVDY_03871 | MagVDY_03870 | Proteobacteria | Alphaproteobacteria |
| Magnetovibrio blakemorei | WP_069958443 | WP_069958444 | WP_069958445 | Proteobacteria | Alphaproteobacteria |
| Mangrovibacter phragmitis | WP_064601340 | WP_064601339 | WP_064601338 | Proteobacteria | Gammaproteobacteria |
| Marasmitruncus massiliensis | WP_101909651 | WP_101909654 | WP_101909655 | Firmicutes | Clostridia |
| Marinilabilia sp. WTE | WP_109262942 | WP_109262939 | WP_109262938 | Bacteroidetes | Bacteroidia |
| Marinobacter sp. ES-1 | WP_022990315 | WP_022990314 | WP_036220149 | Proteobacteria | Gammaproteobacteria |
| Marinobacterium litorale | WP_027854782 | WP_027854783 | WP_027854784 | Proteobacteria | Gammaproteobacteria |
| Mastigocoleus testarum | WP_027846702 | WP_058184423 | WP_027846727 | Cyanobacteria | - |
| Megasphaera cerevisiae | WP_074502485 | WP_074502490 | WP_074502493 | Firmicutes | Negativicutes |
| Methanobacteriales archaeon HGW-Methanobacteriales-1 | PKL66935 | PKL66932 | PKL66931 | Euryarchaeota | Methanobacteria |
| Methanobacterium paludis | WP_013826522 | WP_013826519 | WP_013826518 | Euryarchaeota | Methanobacteria |
| Methanobrevibacter cuticularis | WP_067259993 | WP_067259996 | WP_067259997 | Euryarchaeota | Methanobacteria |
| Methanocaldococcus infernus ME | WP_013099459 | WP_013099456 | WP_013099455 | Euryarchaeota | Methanococci |
| Methanocella arvoryzae | WP_012035713 | WP_012035710 | WP_012035709 | Euryarchaeota | Methanomicrobia |
| Methanococcus vannielii SB | WP_011971883 | WP_011971880 | WP_011971879 | Euryarchaeota | Methanococci |
| Methanolacinia petrolearia DSM 11571 | Mpet_0263 | Mpet_0260 | Mpet_0259 | Euryarchaeota | Methanomicrobia |
| Methanolobus psychrotolerans | WP_094228266 | WP_094228263 | WP_094228262 | Euryarchaeota | Methanomicrobia |
| Methanomassiliicoccus luminyensis | WP_026069118 | WP_019178611 | WP_026069119 | Euryarchaeota | Thermoplasmata |
| Methanosarcina acetivorans C2A | MA3895 | MA3898 | MA3899 | Euryarchaeota | Methanomicrobia |
| Methanosphaerula palustris | WP_012617253 | WP_012617250 | WP_012617249 | Euryarchaeota | Methanomicrobia |
| Methanospirillum stamsii | WP_109942292 | WP_109942295 | WP_109942296 | Euryarchaeota | Methanomicrobia |
| Methanothermobacter marburgensis | WP_013294983 | WP_013294986 | WP_013294987 | Euryarchaeota | Methanobacteria |
| Methanothermococcus thermolithotrophicus | WP_018154785 | WP_018154782 | WP_018154781 | Euryarchaeota | Methanococci |
| Methanothrix soehngenii GP-6 | MCON_0607 | MCON_0610 | MCON_0611 | Euryarchaeota | Methanomicrobia |
| Methanotorris igneus | WP_013799857 | WP_013799860 | WP_013799861 | Euryarchaeota | Methanococci |
| Methylobacterium nodulans ORS 2060 | Mnod_3996 | Mnod_3995 | Mnod_3994 | Proteobacteria | Alphaproteobacteria |
| Methylocaldum szegediense | WP_026610013 | WP_026610014 | WP_026610015 | Proteobacteria | Gammaproteobacteria |
| Methylocystis parvus OBBP | O5ODRAFT_01867 | O5ODRAFT_01868 | O5ODRAFT_01869 | Proteobacteria | Alphaproteobacteria |
| Methylomicrobium buryatense | WP_017842415 | WP_017842414 | WP_017842413 | Proteobacteria | Gammaproteobacteria |
| Methyloversatilis sp. RAC08 | WP_069038340 | WP_069038341 | WP_069038342 | Proteobacteria | Betaproteobacteria |
| Moorella thermoacetica | WP_071521047 | WP_071521044 | WP_071521043 | Firmicutes | Clostridia |
| Myxosarcina sp. GI1 | WP_036489259 | WP_036489262 | WP_081942974 | Cyanobacteria | - |
| Neiella marina | WP_087506928 | WP_087506927 | WP_087506926 | Proteobacteria | Gammaproteobacteria |
| Nitrospirae bacterium GWA2_46_11 | OGW20163 | OGW20164 | OGW20165 | Nitrospirae | - |
| Nitrospirae bacterium GWC2_57_13 | OGW27708 | OGW27709 | OGW27710 | Nitrospirae | - |
| Nodosilinea nodulosa | WP_017298849 | WP_017298848 | WP_017298847 | Cyanobacteria | - |
| Novosphingobium naphthalenivorans | WP_067734406 | WP_067734405 | WP_067734404 | Proteobacteria | Alphaproteobacteria |
| Opitutaceae bacterium EW11 | WP_107745278 | WP_107745279 | WP_107745280 | Verrucomicrobia | Opitutae |
| Opitutaceae bacterium TSB47 | AW736_03595 | AW736_03600 | AW736_03605 | Verrucomicrobia | Verrucomicrobia |
| Orenia marismortui | WP_018247481 | WP_018247478 | WP_018247477 | Firmicutes | Clostridia |
| Oscillatoria sp. PCC 10802 | WP_017716686 | WP_017716685 | WP_017716684 | Cyanobacteria | - |
| Oscillochloris trichoides DG-6 | EFO82066 | EFO82064 | EFO82063 | Chloroflexi | Chloroflexia |
| Oxobacter pfennigii | WP_054874024 | WP_054874027 | WP_054874028 | Firmicutes | Clostridia |
| Paenibacillus stellifer DSM 14472 | Ga0069362_114164 | Ga0069362_114163 | Ga0069362_114162 | Firmicutes | Bacilli |
| Paludibacter propionicigenes | WP_013444938 | WP_013444941 | WP_013444942 | Bacteroidetes | Bacteroidia |
| Paludibacterium yongneupense | WP_028534981 | WP_028534980 | WP_028534979 | Proteobacteria | Betaproteobacteria |
| Pantoea cypripedii LMG 2657 | Ga0309980_12282 | Ga0309980_12281 | Ga0309980_12280 | Proteobacteria | Gammaproteobacteria |
| Pararhodospirillum photometricum DSM 122 | RSPPHO_02584 | RSPPHO_02583 | RSPPHO_02582 | Proteobacteria | Alphaproteobacteria |
| Pelobacter propionicus DSM 2379 | Ppro_3467 | Ppro_3468 | Ppro_3469 | Proteobacteria | Deltaproteobacteria |
| Pelosinus propionicus DSM 13327 | Ga0074834_1010107 | Ga0074834_1010104 | Ga0074834_1010103 | Firmicutes | Negativicutes |
| Peptococcaceae bacterium BRH_c4a | KJR99751 | KJR99748 | KJR99747 | Firmicutes | Clostridia |
| Peptococcaceae bacterium BRH_c8a | KJS11410 | KJS11413 | KJS11414 | Firmicutes | Clostridia |
| Peptococcaceae bacterium CEB3 | WP_047829845 | WP_047829846 | WP_047829847 | Firmicutes | Clostridia |
| Petroclostridium xylanilyticum | WP_094550765 | WP_094550759 | WP_094550757 | Firmicutes | Clostridia |
| Phaeospirillum fulvum DSM 114 | Ga0061132_00876 | Ga0061132_00875 | Ga0061132_00874 | Proteobacteria | Alphaproteobacteria |
| Phormidesmis priestleyi Ana | KPQ34319 | KPQ34320 | KPQ34321 | Cyanobacteria | - |
| Pleomorphomonas sp. 86 | Ga0302604_11600 | Ga0302604_11601 | Ga0302604_11602 | Proteobacteria | Alphaproteobacteria |
| Prevotella oryzae | WP_036881131 | WP_036880219 | WP_036880217 | Bacteroidetes | Bacteroidia |
| Propionispira arboris | WP_091831461 | WP_091831467 | WP_091831469 | Firmicutes | Negativicutes |
| Propionispora vibrioides DSM 13305 | Ga0074833_1194 | Ga0074833_1197 | Ga0074833_1198 | Firmicutes | Negativicutes |
| Pseudanabaena sp. SR411 | WP_094531215 | WP_094531213 | WP_094531211 | Cyanobacteria | - |
| Pseudodesulfovibrio profundus | WP_097013057 | WP_097013060 | WP_097013061 | Proteobacteria | Deltaproteobacteria |
| Pseudomonas stutzeri A1501 | PST_1326 | PST_1327 | PST_1328 | Proteobacteria | Gammaproteobacteria |
| Rahnella sp. AA | WP_101076633 | WP_101076632 | WP_101076631 | Proteobacteria | Gammaproteobacteria |
| Raoultella ornithinolytica JUb54 | Ga0304837_101468 | Ga0304837_101467 | Ga0304837_101466 | Proteobacteria | Gammaproteobacteria |
| Rhodobacter capsulatus A52 | Ga0100862_105159 | Ga0100862_105160 | Ga0100862_105161 | Proteobacteria | Alphaproteobacteria |
| Rhodoblastus acidophilus DSM 137 | Ga0170454_10611 | Ga0170454_10612 | Ga0170454_10613 | Proteobacteria | Alphaproteobacteria |
| Rhodomicrobium vannielii ATCC 17100 | Rvan_1953 | Rvan_1954 | Rvan_1955 | Proteobacteria | Alphaproteobacteria |
| Rhodopila globiformis | WP_104518213 | WP_104518212 | WP_104518211 | Proteobacteria | Alphaproteobacteria |
| Rhodopseudomonas palustris CGA009 | RPA4620 | RPA4619 | RPA4618 | Proteobacteria | Alphaproteobacteria |
| Rhodospirillum rubrum ATCC 11170 | Rru_A1010 | Rru_A1011 | Rru_A1012 | Proteobacteria | Alphaproteobacteria |
| Rhodovibrio salinarum | WP_027289758 | WP_027289759 | WP_027289760 | Proteobacteria | Alphaproteobacteria |
| Rhodovulum sp. PH10 | A33M_1296 | A33M_1297 | A33M_1298 | Proteobacteria | Alphaproteobacteria |
| Richelia intracellularis HH01 | CCH66462 | CCH66469 | CCH66470 | Cyanobacteria | - |
| Roseiflexus castenholzii | WP_012122497 | WP_012122495 | WP_012122494 | Chloroflexi | Chloroflexia |
| Roseospirillum parvum | WP_092620711 | WP_092620709 | WP_092620707 | Proteobacteria | Alphaproteobacteria |
| Saccharicrinis fermentans | WP_027473409 | WP_027473406 | WP_027473405 | Bacteroidetes | Bacteroidia |
| Sediminispirochaeta smaragdinae | WP_013255569 | WP_013255566 | WP_013255565 | Spirochaetes | Spirochaetia |
| Smithella sp. SCADC | KFO68400 | KFO68403 | KFO68404 | Proteobacteria | Deltaproteobacteria |
| Spirochaeta cellobiosiphila | WP_028973004 | WP_028973001 | WP_028973000 | Spirochaetes | Spirochaetia |
| Spirochaeta thermophila | WP_013313167 | WP_013313164 | WP_013313163 | Spirochaetes | Spirochaetia |
| Spirochaetes bacterium GWB1_27_13 | OHD05375 | OHD05378 | OHD05379 | Spirochaetes | - |
| Spirochaetes bacterium GWB1_36_13 | OHD11363 | OHD11360 | OHD11367 | Spirochaetes | - |
| Spirochaetes bacterium GWE1_32_154 | OHD39357 | OHD39354 | OHD39353 | Spirochaetes | - |
| Sporolactobacillus terrae | WP_028984127 | WP_028984128 | WP_028984129 | Firmicutes | Bacilli |
| Sporomusa acidovorans | WP_093792115 | WP_093792109 | WP_093792107 | Firmicutes | Negativicutes |
| Sulfuricurvum kujiense YK-1, DSM 16994 | Sulku_1500 | Sulku_1498 | Sulku_1497 | Proteobacteria | Epsilonproteobacteria |
| Sulfuriferula sp. AH1 | WP_087445781 | WP_087445782 | WP_087445783 | Proteobacteria | Betaproteobacteria |
| Sulfurimonas sp. | PNV83262 | PNV83260 | PNV83259 | Proteobacteria | Epsilonproteobacteria |
| Sulfurospirillum cavolei | WP_060826056 | WP_060826054 | WP_082709164 | Proteobacteria | Epsilonproteobacteria |
| Sulfurovum sp. enrichment culture clone C5 | CUV65475 | CUV65477 | CUV65478 | Proteobacteria | Epsilonproteobacteria |
| Syntrophobacter fumaroxidans MPOB | WP_011697884 | WP_011697881 | WP_011697880 | Proteobacteria | Deltaproteobacteria |
| Syntrophobotulus glycolicus FlGlyR, DSM 8271 | Sgly_2841 | Sgly_2840 | Sgly_2839 | Firmicutes | Clostridia |
| Syntrophomonas zehnderi | WP_046497554 | WP_046497545 | WP_046497542 | Firmicutes | Clostridia |
| Syntrophothermus lipocalidus DSM 12680 | WP_013176275 | WP_013176272 | WP_013176271 | Firmicutes | Clostridia |
| Telmatospirillum siberiense | WP_101250570 | WP_101250571 | WP_101250572 | Proteobacteria | Alphaproteobacteria |
| Teredinibacter turnerae | WP_018277248 | WP_018277247 | WP_018277246 | Proteobacteria | Gammaproteobacteria |
| Terrimicrobium sacchariphilum | WP_075078689 | WP_084400275 | WP_075078690 | Verrucomicrobia | Spartobacteria |
| Thauera sp. D20 | WP_107492233 | WP_107492234 | WP_107492235 | Proteobacteria | Betaproteobacteria |
| Thermacetogenium phaeum | WP_015049976 | WP_015049979 | WP_015049980 | Firmicutes | Clostridia |
| Thermicanus aegyptius | WP_028986272 | WP_028986273 | WP_028986274 | Firmicutes | Bacilli |
| Thermincola potens JR | TherJR_0688 | TherJR_0691 | TherJR_0692 | Firmicutes | Clostridia |
| Thermoanaerobacterium thermosaccharolyticum | WP_013298321 | WP_013298320 | WP_013298319 | Firmicutes | Clostridia |
| Thermodesulfovibrio yellowstonii DSM 11347 | WP_012546613 | WP_012546033 | WP_012545394 | Nitrospirae | Nitrospira |
| Thioflexothrix psekupsii | WP_086487138 | WP_086487137 | WP_086487136 | Proteobacteria | Gammaproteobacteria |
| Thiohalocapsa sp. ML1 | WP_058554369 | WP_058554368 | WP_058554367 | Proteobacteria | Gammaproteobacteria |
| Thiorhodococcus drewsii AZ1 | ThidrDRAFT_2681 | ThidrDRAFT_2682 | ThidrDRAFT_2683 | Proteobacteria | Gammaproteobacteria |
| Thiothrix nivea DSM 5205 | EIJ36507 | EIJ36508 | EIJ36509 | Proteobacteria | Gammaproteobacteria |
| Tolumonas lignilytica BRL6-1 | H027DRAFT2334 | H027DRAFT2335 | H027DRAFT2336 | Proteobacteria | Gammaproteobacteria |
| Treponema primitia | WP_010260142 | WP_010260134 | WP_010260131 | Spirochaetes | Spirochaetia |
| Trichodesmium erythraeum | WP_011613474 | WP_011613475 | WP_011613476 | Cyanobacteria |  |
| Uliginosibacterium gangwonense | WP_018607902 | WP_018607903 | WP_018607904 | Proteobacteria | Betaproteobacteria |
| uncultured Alphaproteobacteria bacterium | SBW06912 | SBW06906 | SBW06899 | Proteobacteria | Alphaproteobacteria |
| unicellular cyanobacterium SU2 | WP_085434912 | WP_085434911 | WP_085434910 | Cyanobacteria | - |
| Verrucomicrobia bacterium LP2A | WP_024807527 | WP_024807526 | WP_024807525 | Verrucomicrobia | - |
| Verrucomicrobia bacterium Tous-C9LFEB | PAW77340 | PAW77337 | PAW77336 | Verrucomicrobia | - |
| Verrucomicrobiae bacterium DG1235 | WP_040901465 | WP_008103118 | WP_040899453 | Verrucomicrobia | Verrucomicrobiae |
| Vibrio diazotrophicus | WP_102939852 | WP_102939851 | WP_102939850 | Proteobacteria | Gammaproteobacteria |
| Vulcanococcus limneticus | WP_094591659 | WP_094591660 | WP_094591661 | Cyanobacteria |  |
| Xanthobacter autotrophicus Py2 | Xaut_0088 | Xaut_0089 | Xaut_0090 | Proteobacteria | Alphaproteobacteria |
| Yangia sp. CCB-MM3 | WP_066099558 | WP_066099560 | WP_066099562 | Proteobacteria | Alphaproteobacteria |
| Youngiibacter fragilis | WP_023388904 | WP_023388907 | WP_023388908 | Firmicutes | Clostridia |
| Zavarzinia sp. HR-AS | WP_109903383 | WP_109903385 | WP_109903387 | Proteobacteria | Alphaproteobacteria |
| Zetaproteobacteria bacterium CG17_big_fil_post_rev_8_21_14_2_50_50_13 | PIQ34264 | PIQ34263 | PIQ34262 | Proteobacteria | Zetaproteobacteria |
| Zoogloea sp. LCSB751 | WP_079432933 | WP_079432932 | WP_079432931 | Proteobacteria | Betaproteobacteria |
| Zymomonas mobilis | WP_011241556 | WP_011241557 | WP_011241558 | Proteobacteria | Alphaproteobacteria |
| **Species** | **AnfH** | **AnfD** | **AnfK** | **Phylum** | **Class** |
| Azotobacter vinelandii DJ, ATCC BAA-1303 | Avin_49000 | Avin_48990 | Avin_48970 | Proteobacteria | Gammaproteobacteria |
| Bacteroidales bacterium Barb6 | WP_066182251 | WP_066186490 | WP_066182235 | Bacteroidetes | Bacteroidia |
| Brenneria salicis ATCC 15712 | Bresa_00753 | Bresa_00754 | Bresa_00756 | Proteobacteria | Gammaproteobacteria |
| Chloroherpeton thalassium ATCC 35110 | Ctha_1828 | Ctha_1831 | Ctha_1833 | Chlorobi | Chlorobia |
| Clostridium pasteurianum DSM 525, ATCC 6013 | Ga0069491_113885 | Ga0069491_113883 | Ga0069491_113881 | Firmicutes | Clostridia |
| Dendrosporobacter quercicolus | WP_092072016 | WP_092072026 | WP_092072032 | Firmicutes | Negativicutes |
| Desulfitobacterium chlororespirans | WP_072772837 | WP_072772834 | WP_072772832 | Firmicutes | Clostridia |
| Desulfovibrio sp. 86-1 | Ga0302701_111207 | Ga0302701_111210 | Ga0302701_111212 | Proteobacteria | Deltaproteobacteria |
| Dysgonomonas capnocytophagoides | WP_026625686 | WP_026625683 | WP_026625681 | Bacteroidetes | Bacteroidia |
| Opitutaceae bacterium TSB47 | AW736_03515 | AW736_03520 | AW736_03530 | Verrucomicrobia | Verrucomicrobia |
| Pantoea cypripedii LMG 2657 | Ga0309980_12203 | Ga0309980_12204 | Ga0309980_12206 | Proteobacteria | Gammaproteobacteria |
| Pararhodospirillum photometricum DSM 122 | RSPPHO_00490 | RSPPHO_00489 | RSPPHO_00487 | Proteobacteria | Alphaproteobacteria |
| Rhodobacter capsulatus A52 | Ga0100862_105146 | Ga0100862_105145 | Ga0100862_105143 | Proteobacteria | Alphaproteobacteria |
| Rhodoblastus acidophilus DSM 137 | Ga0170454_11380 | Ga0170454_11381 | Ga0170454_11383 | Proteobacteria | Alphaproteobacteria |
| **Species** | **VnfH** | **VnfD** | **VnfK** | **Phylum** | **Class** |
| Azomonas agilis DSM 375 | LX59DRAFT_02210 | LX59DRAFT_01075 | LX59DRAFT_01073 | Proteobacteria | Gammaproteobacteria |
| Azotobacter vinelandii DJ, ATCC BAA-1303 | Avin_02660 | Avin_02610 | Avin_02590 | Proteobacteria | Gammaproteobacteria |
| Chromatiales bacterium | PHBDraft_25530 | PHBDraft_25580 | PHBDraft_25610 | Proteobacteria | Gammaproteobacteria |
| Clostridium kluyveri DSM 555 | CKL_1748 | CKL_1747 | CKL_1745 | Firmicutes | Clostridia |
| Desulfobacter curvatus DSM 3379 | B147DRAFT_02029 | B147DRAFT_02028 | B147DRAFT_02026 | Proteobacteria | Deltaproteobacteria |
| Ethanoligenens harbinense YUAN-3T, DSM 18485 | Ethha_2309 | Ethha_2312 | Ethha_2314 | Firmicutes | Clostridia |
| Magnetospirillum bellicus VDY | MagVDY_01910 | MagVDY_01908 | MagVDY_01906 | Proteobacteria | Alphaproteobacteria |
| Methanosarcina acetivorans C2A | MA1213 | MA1216 | MA1218 | Euryarchaeota | Methanomicrobia |
| Methylocystis parvus OBBP | O5ODRAFT_02619 | O5ODRAFT_02621 | O5ODRAFT_02623 | Proteobacteria | Alphaproteobacteria |
| Paenibacillus durus P3L-5 | L664DRAFT_00627 | L664DRAFT_00626 | L664DRAFT_00624 | Firmicutes | Bacilli |
| Phaeospirillum fulvum DSM 114 | Ga0061132_02947 | Ga0061132_02945 | Ga0061132_02943 | Proteobacteria | Alphaproteobacteria |
| Rhodoblastus acidophilus DSM 137 | Ga0170454_103245 | Ga0170454_103247 | Ga0170454_103249 | Proteobacteria | Alphaproteobacteria |
| Rhodopseudomonas palustris CGA009 | RPA1376 | RPA1378 | RPA1380 | Proteobacteria | Alphaproteobacteria |
| Tolumonas lignilytica BRL6-1 | H027DRAFT2934 | H027DRAFT2940 | H027DRAFT2942 | Proteobacteria | Gammaproteobacteria |
| **Species** | **Bch/ChlL** | **Bch/ChlN** | **Bch/ChlB** | **Phylum** | **Class** |
| Chlorobium phaeobacteroides DSM 266 | WP_011746157 | WP_011746159 | WP_011746158 | Chlorobi | Chlorobia |
| Erythrobacter sp. NAP1 | WP_007165005 | WP_007165002 | WP_007165003 | Proteobacteria | Alphaproteobacteria |
| Gloeobacter violaceus PCC 7421 | WP_011142366 | WP_011142365 | WP_011140219 | Cyanobacteria | Gloeobacteria |
| Halorhodospira halophila | WP_011814420 | WP_011814423 | WP_011814422 | Proteobacteria | Gammaproteobacteria |
| Heliobacterium modesticaldum | WP_012282383 | WP_012282384 | WP_012282385 | Firmicutes | Clostridia |
| Hoeflea phototrophica DFL-43 | WP_040449113 | WP_007196612 | WP_007196611 | Proteobacteria | Alphaproteobacteria |
| Roseiflexus castenholzii | WP_012120059 | WP_012120057 | WP_012120058 | Chloroflexi | Chloroflexia |
| Synechococcus sp. CC9311 | WP_011619890 | WP_011619888 | WP_011619889 | Cyanobacteria | - |
| Synechococcus sp. JA-2-3B'a(2-13) | WP_011432649 | WP_011432647 | WP_011433773 | Cyanobacteria | - |
| Synechococcus sp. RCC307 | WP_011935983 | WP_011935981 | WP_011935982 | Cyanobacteria | - |
