## Supplemental File 2 for "Reconstructing the Evolutionary History of Nitrogenases: Evidence for Ancestral Molybdenum-Cofactor Utilization"

**Expanded nitrogenase phylogenetic analyses**

*AncE* *node selection across alternate phylogenetic topologies*

Five nitrogenase phylogenies were reconstructed by varying alignment methods and amino acid substitution and rate heterogeneity models (Tree-1–5; Figure S1). The topologies across the five trees are largely similar excepting root position. Due to these differences in root position, equivalent AncE nodes to not exist for all five trees. In Tree-1, Tree-2, and Tree-4, AncE is designated as the node ancestral to Nif-I and Nif-II clades. However, in Tree-3 and Tree-5, AncE is instead designated as the node ancestral to Nif-II and uncharacterized/Vnf/Anf clades. These AncE nodes would otherwise be equivalent if the trees were unrooted.

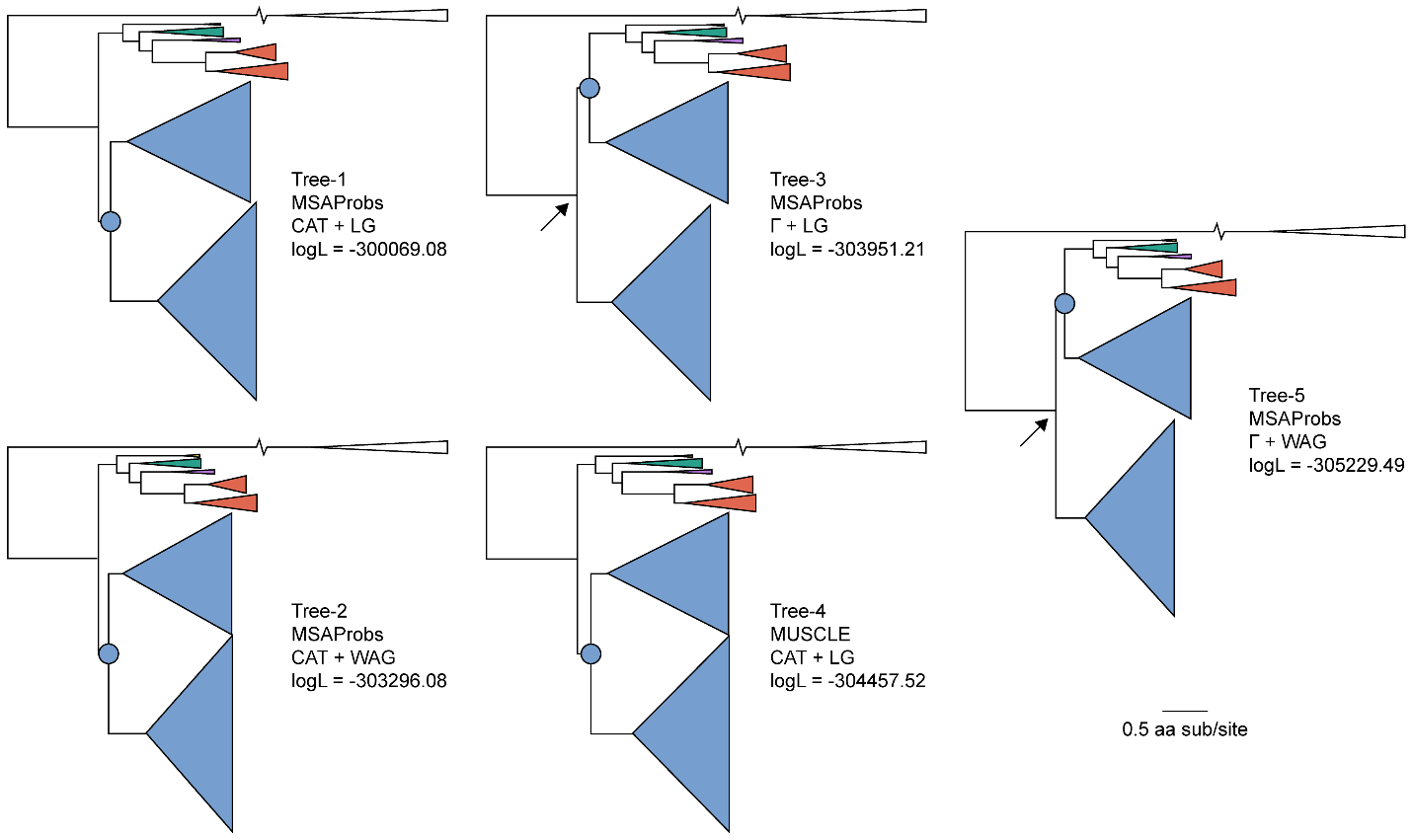

**Figure S1.** Top five maximum likelihood nitrogenase phylogenies, ranked by log likelihood values. Arrow indicates root position difference in Tree-3 and Tree-5 as compared to the Tree-1, Tree-2, and Tree-4. AncE node selection differs across different topologies, indicated by a blue circle. Branch length scale is in units of amino acid substitutions per site. Phylogeny coloring is as follows: Clfx, yellow; F-Mc, green; Mb-Mc, purple; Anf/Vnf, red; Nif-I/-II, blue.

In order to assess the sensitivity of AncE ancestral sequence composition to root position differences, we calculated total HDK sequence identities of AncE across all alternate topologies (Tree1–5; Figure S2). Sequence identity across all AncE sequences ranges between ~89 and 99%. The sequence identity between AncE-1 and AncE-3, derived from topologies with different root positions, is ~99%. By comparison, the sequence identity between AncE-1 and AncE-4, derived from topologies with the same root positions but reconstructed from a different sequence alignment method, is ~91%. AncE-4, the only sequence derived from a tree reconstructed from a MUSCLE alignment, consistently has the lowest identity with other AncE sequences (~89–91%), the latter of which are all derived from trees reconstructed from a MSAProbs alignment. Therefore, we do not find evidence that root position differences affect AncE sequence composition more than the sequence alignment for phylogenetic reconstruction. The analogous node designations for AncE ancestors across alternate topologies is thus appropriate for subsequent sequence analysis.

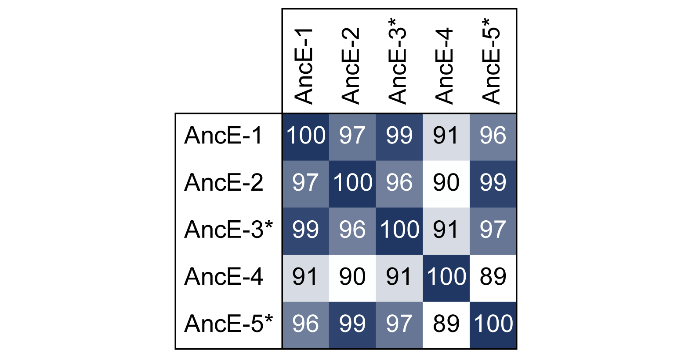

**Figure S2.** AncE total HDK sequence identities. Asterisk (*) denotes AncE sequences derived from phylogenies (Tree-3, Tree-5) with differing root position than the most likely phylogeny, Tree-1.

*Nitrogenase tree topology with an expanded outgroup*

Due to differences in root position among the five reconstructed nitrogenase phylogenies, we performed an additional reconstruction with an expanded outgroup. The outgroup was expanded from 10 light-independent protochlorophyllide reductase (Bch/ChlLNB) sequences to 323 sequences alongside the 284-sequence nitrogenase dataset. Bch/ChlLNB homologs were identified by BLASTp using query sequences from *Chlorobium phaeobacteroides* (BchL, WP_011746157; BchN, WP_011746159; BchB, WP_011746158) with an Expect value cutoff of <1e-5. All sequences were realigned using MAFFT^1^ and alignments were trimmed using TrimAl^2^ (“gappyout” option). Trees were reconstructed using RAxML^3^ under the LG model of protein sequence evolution^4^ and either the gamma distribution model of rate heterogeneity^5^ or its approximation (CAT^6^), with branch support calculated by rapid bootstrapping. In sum, two trees were reconstructed with an unexpanded outgroup, and two trees with an expanded outgroup. The CAT tree branch lengths and model parameters were then reoptimized under the gamma model to make likelihoods comparable. Tree topologies were compared using Shimodaira’s Approximately Unbiased (AU) test^7^ implemented in CONSEL^8^, where a topology can be rejected if p<0.05.

For the two phylogenies reconstructed with an expanded outgroup, branching order of the major clades does not differ between those reconstructed by the gamma rate heterogeneity model and the CAT approximation, and is the same as that for Tree-1, Tree-2, and Tree-4 (Table S2, Figure S1). The root position is supported in both expanded-outgroup trees with a bootstrap value of 100, as compared to values of 56 and 25 for the two unexpanded-outgroup trees. stabilizing effect of outgroup expansion on tree topology lends support to the topology consistently recovered after outgroup expansion (equivalent to Tree-1, -2, and -4), relative to alternate topologies recovered under model approximations with a smaller outgroup (Trees-3 and -5). However, alternate topologies were not rejected by the AU test (Table S2), and as such can be retained as plausible hypotheses for evolutionary relationships among nitrogenase proteins.

**Table S2**. Log likelihood values and AU test p-values for nitrogenase phylogenies reconstructed with an expanded outgroup.

| Tree | Log likelihood | | AU test p-value | |
| --- | --- | --- | --- | --- |
|  | CAT | Γ | CAT | Γ |
| 10-sequence outgroup | -298569.41 | -298569.70 | 0.383 | 0.617 |
| 323-sequence outgroup | -327157.24 | -327148.94 | 0.456 | 0.544 |
