## Supplemental File 3 for "Reconstructing the Evolutionary History of Nitrogenases: Evidence for Ancestral Molybdenum-Cofactor Utilization"

**Table S3.** Posterior probabilities of maximum likelihood nitrogenases, averaged across total HDK sequence and across 30 active-site residues, defined as those within 5 Å of any atom within the FeMo-cofactor (PDB 3U7Q^1^) or FeV-cofactor (PDB 5N6Y^2^).

| Ancestor | Mean ancestral site posterior probability (± 1σ) | Mean ancestral active-site posterior probability (± 1σ) |
| --- | --- | --- |
| AncA-1 | 0.90 (± 0.18) | 0.93 (± 0.16) |
| AncA-2 | 0.91 (± 0.17) | 0.92 (± 0.19) |
| AncA-3 | 0.90 (± 0.18) | 0.94 (± 0.14) |
| AncA-4 | 0.89 (± 0.19) | 0.94 (± 0.14) |
| AncA-5 | 0.91 (± 0.17) | 0.92 (± 0.19) |
| AncB-1 | 0.86 (± 0.21) | 0.94 (± 0.15) |
| AncB-2 | 0.87 (± 0.20) | 0.94 (± 0.14) |
| AncB-3 | 0.86 (± 0.21) | 0.93 (± 0.16) |
| AncB-4 | 0.85 (± 0.22) | 0.94 (± 0.15) |
| AncB-5 | 0.87 (± 0.20) | 0.94 (± 0.14) |
| AncC-1 | 0.88 (± 0.20) | 0.97 (± 0.10) |
| AncC-2 | 0.89 (± 0.19) | 0.98 (± 0.07) |
| AncC-3 | 0.88 (± 0.20) | 0.96 (± 0.12) |
| AncC-4 | 0.87 (± 0.21) | 0.96 (± 0.12) |
| AncC-5 | 0.89 (± 0.19) | 0.98 (± 0.07) |
| AncD-1 | 0.84 (± 0.22) | 0.96 (± 0.12) |
| AncD-2 | 0.85 (± 0.21) | 0.97 (± 0.09) |
| AncD-3 | 0.84 (± 0.22) | 0.95 (± 0.13) |
| AncD-4 | 0.83 (± 0.23) | 0.96 (± 0.10) |
| AncD-5 | 0.86 (± 0.21) | 0.97 (± 0.09) |
| AncE-1 | 0.86 (± 0.21) | 0.94 (± 0.15) |
| AncE-2 | 0.87 (± 0.21) | 0.94 (± 0.15) |
| AncE-3 | 0.86 (± 0.21) | 0.94 (± 0.15) |
| AncE-4 | 0.86 (± 0.21) | 0.94 (± 0.15) |
| AncE-5 | 0.87 (± 0.20) | 0.94 (± 0.16) |

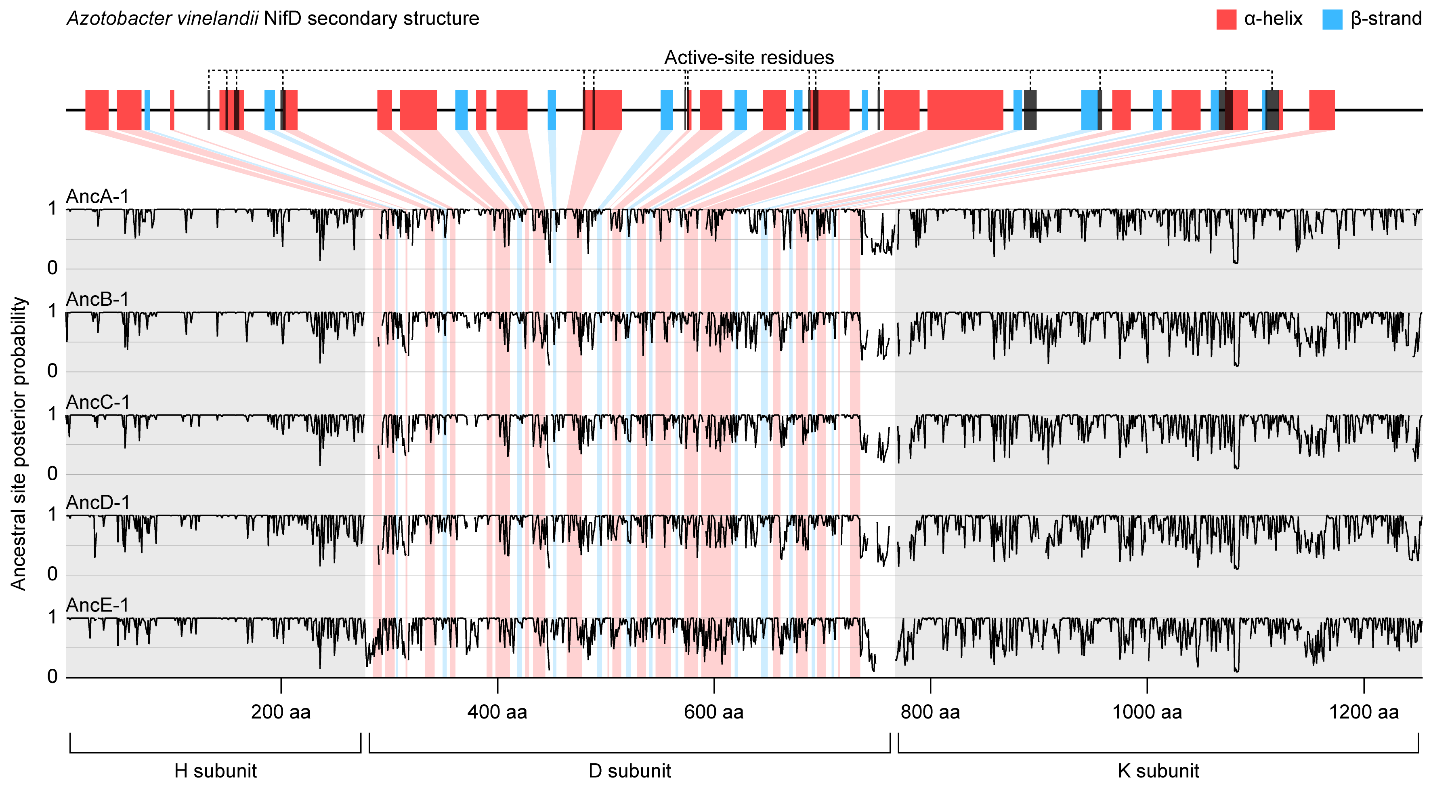

**Figure S4.** Ancestral site posterior probabilities of aligned ancestral nitrogenase sequences inferred from the most likely reconstructed phylogeny (Tree-1). The aligned secondary structure of the *A. vinelandii* NifD subunit is shown on top (from PDB 3U7Q^16^). Active-site residues include 30 amino acids positioned within 5 Å of the active-site cofactor.
