## Supplemental File 4 for "Reconstructing the Evolutionary History of Nitrogenases: Evidence for Ancestral Molybdenum-Cofactor Utilization"

**Table S4.** Nitrogenase active-site residues uniquely conserved in each clade or group of clades of known metal dependence. Each unique residue is listed in the same row as the homologous residue of *A. vinelandii*. Below each unique residue are uncharacterized or ancestral sequences that share that residue. All residue numbering based on *A. vinelandii* NifD. All residues are within 5 Å of any atom of the FeMo-cofactor (PDB 37UQ^1^) or FeV-cofactor (PDB 5N6Y^2^).

| *A. vinelandii* active-site residue | Homologous unique residues | | | |
| --- | --- | --- | --- | --- |
|  | Anf | Vnf | Anf/Vnf | Nif-I/-II |
| Ile-59 |  |  |  |  |
| Ala-65 |  |  | Cys-65  ^AncA^ | Ala-65  ^Clfx, F-Mc, Mb-Mc,^ ^AncB–E^ |
| Gly-69 | His-69  ^Anc-A^ | Leu-69 |  |  |
| Val-70 |  |  |  |  |
| Gly-95 |  |  |  |  |
| Arg-96 |  |  | Lys-96  ^Mb-Mc, AncA–B^ | Arg-96  ^Clfx, F-Mc, AncC–E^ |
| Gln-191 |  |  |  |  |
| His-195 |  |  |  |  |
| Tyr-229 |  |  |  |  |
| Ile-231 |  |  |  |  |
| Cys-275 |  |  |  |  |
| Arg-277 |  |  |  |  |
| Ser-278 |  |  |  |  |
| Phe-300 |  | Trp-300  ^Clfx^ |  |  |
| Ile-355 | Pro-355 | Thr-355  ^AncA–B^ |  |  |
| Gly-356 |  |  |  |  |
| Gly-357 |  |  |  |  |
| Leu-358 |  | Pro-358  ^Clfx, F-Mc, Mb-Mc,^ ^AncA–D^ |  |  |
| Arg-359 | Lys-359  ^Mb-Mc^ |  |  |  |
| Pro-360 |  |  | Leu-360  ^AncA^ |  |
| Glu-380 |  |  | Lys-380  ^AncA^ |  |
| Phe-381 |  |  |  |  |
| Gly-424 |  |  |  |  |
| Ile-425 |  | Pro-425  ^AncA-3–4^ |  |  |
| Lys-426 |  |  | Arg-426  ^AncA^ |  |
| Glu-427 | Pro-427 | Val-427  ^AncA^ |  |  |
| Gln-440 |  |  | Asn-440  ^Clfx, F-Mc, AncA–D^ | Gln-440  ^AncE^ |
| Met-441 | Ala-441  ^AncA-1, AncA-3–4^ |  |  |  |
| His-442 |  |  |  |  |
| Ser-443 |  |  |  |  |
