## Supplemental File 5 for "Reconstructing the Evolutionary History of Nitrogenases: Evidence for Ancestral Molybdenum-Cofactor Utilization"

**Expanded nitrogenase pocket volume analyses**

Structural homology modeling of 33 extant and ancestral (25 maximum likelihood and 2,500 Bayesian-sampled) nitrogenase D-subunit proteins was performed using 38 NifD and 2 VnfD structural templates (Table S5). Two separate modeling analyses were performed in order to assess the effect of cofactor type on the calculated pocket volume of generated models: all nitrogenase sequences were modeled by specifying the inclusion of (1) the FeMo-cofactor of the 3U7Q NifD structure^1^ or (2) the FeV-cofactor of the 56NY VnfD template^2^. Active-site pocket volumes were calculated for both FeMo-cofactor- and FeV-cofactor-modelled structures. For each cofactor analysis, we grouped pocket volume data by nitrogenase sequence and performed a Kruskal-Wallis test, which tests whether at least one group differs significantly from other groups, followed by Dunn’s test, which tests for median difference in pairwise comparisons of groups. All statistical tests were performed in R.

**Table S5.** Structural templates used for homology modeling of ancestral and extant nitrogenase D-subunit sequences. Pocket volume calculated for published structures by the same manner as for ancestral and extant structural models.

| Protein Data Bank template structure | Active-site cofactor | Organism | Pocket volume (Å^3^) |
| --- | --- | --- | --- |
| 1FP4^3^ | FeMo | *Azotobacter vinelandii* | 963.250 |
| 1G20^4^ | FeMo | *Azotobacter vinelandii* | 966.750 |
| 1G21^4^ | FeMo | *Azotobacter vinelandii* | 1000.000 |
| 1L5H^5^ | FeMo | *Azotobacter vinelandii* | 1246.375 |
| 1M1N^6^ | FeMo | *Azotobacter vinelandii* | 1012.125 |
| 1M1Y^7^ | FeMo | *Azotobacter vinelandii* | 974.125 |
| 1M34^7^ | FeMo | *Azotobacter vinelandii* | 1018.500 |
| 1N2C^8^ | FeMo | *Azotobacter vinelandii* | 962.000 |
| 2AFH^9^ | FeMo | *Azotobacter vinelandii* | 995.125 |
| 2AFI^9^ | FeMo | *Azotobacter vinelandii* | 995.125 |
| 2MIN^10^ | FeMo | *Azotobacter vinelandii* | 981.875 |
| 3K1A^11^ | FeMo | *Azotobacter vinelandii* | 964.125 |
| 3MIN^10^ | FeMo | *Azotobacter vinelandii* | 977.000 |
| 3U7Q^1^ | FeMo | *Azotobacter vinelandii* | 994.750 |
| 4ND8^1^ | FeMo | *Azotobacter vinelandii* | 1009.250 |
| 4TKU^12^ | FeMo | *Azotobacter vinelandii* | 989.125 |
| 4TKV^12^ | FeMo | *Azotobacter vinelandii* | 980.500 |
| 4WNA^13^ | FeMo | *Azotobacter vinelandii* | 985.500 |
| 4WZA^14^ | FeMo | *Azotobacter vinelandii* | 997.250 |
| 4WZB^9^ | FeMo | *Azotobacter vinelandii* | 1015.625 |
| 4XPI^15^ | FeMo | *Azotobacter vinelandii* | 999.750 |
| 5BVG^16^ | FeMo | *Azotobacter vinelandii* | 996.500 |
| 5BVH^16^ | FeMo | *Azotobacter vinelandii* | 994.250 |
| 5CX1^17^ | FeMo | *Azotobacter vinelandii* | 994.125 |
| 5VQ4^18^ | FeMo | *Azotobacter vinelandii* | 995.750 |
| 6BBL^19^ | FeMo | *Azotobacter vinelandii* | 985.125 |
| 6CDK^20^ | FeMo | *Azotobacter vinelandii* | 986.500 |
| 1MIO^21^ | FeMo | *Clostridium pasteuranium* | 1015.250 |
| 4WES^22^ | FeMo | *Clostridium pasteuranium* | 1005.000 |
| 4WN9^13^ | FeMo | *Clostridium pasteuranium* | 1018.125 |
| 5VPW^23^ | FeMo | *Clostridium pasteuranium* | 1073.375 |
| 5VQ3^18^ | FeMo | *Clostridium pasteuranium* | 1023.250 |
| 5KOH^23^ | FeMo | *Gluconacetobacter diazotrophicus* | 963.875 |
| 5KOJ^23^ | FeMo | *Gluconacetobacter diazotrophicus* | 961.750 |
| 1H1L^24^ | FeMo | *Klebsiella pneumoniae* | 1009.000 |
| 1QGU^25^ | FeMo | *Klebsiella pneumoniae* | 998.250 |
| 1QH1^25^ | FeMo | *Klebsiella pneumoniae* | 1000.500 |
| 1QH8^25^ | FeMo | *Klebsiella pneumoniae* | 982.375 |
| 5N6Y^2^ | FeV | *Azotobacter vinelandii* | 938.125 |
| 6FEA^26^ | FeV | *Azotobacter vinelandii* | 948.125 |

Active-site pocket volumes of modeled extant and ancestral nitrogenase are plotted in Figure S4. In general, structures modeled with a FeV-cofactor have larger pocket volumes (mean ≈ 1158.361 Å^3^) than those of structures modeled with a FeMo-cofactor (mean ≈ 1120.920 Å^3^). Despite these bulk differences between FeMo-cofactor and FeV-cofactor-modelled structures, the relative pocket volume trends across major nitrogenase extant clades and ancestors are maintained in both analyses. From the Kruskal-Wallis test, we conclude that at least one pocket volume group differs significantly in both FeMo- and Fe-V- cofactor datasets (*p* < .001). The results of the pairwise comparisons of Dunn’s test are displayed in Figure S5, where we conclude statistically significant median difference for *p* < .025, and median similarity for *p* > .025. We find that, for both FeMo-cofactor and FeV-cofactor analyses, more pairwise comparisons of Vnf versus Nif-II models (derived from nitrogenases with different metal dependence) indicate median pocket volume similarity than pairwise comparisons of Nif-I versus Nif-II models (derived from nitrogenases with the same metal dependence). We also find that, in both analyses, most ancestral models have pocket volume median similarity to both Vnf and Nif-II models, with the exception of AncA models, which have similar pocket volume median similarity to both Vnf and Anf models. The results of Dunn’s test therefore suggest that pocket volume cannot reliably predict nitrogenase metal dependence, and that pocket volume cannot unambiguously be used to classify the metal dependence of ancestral nitrogenases. These findings apply to both FeMo-cofactor and FeV-cofactor analyses, as well as to both maximum likelihood and Bayesian-sampled ancestral analyses (Figure S6).

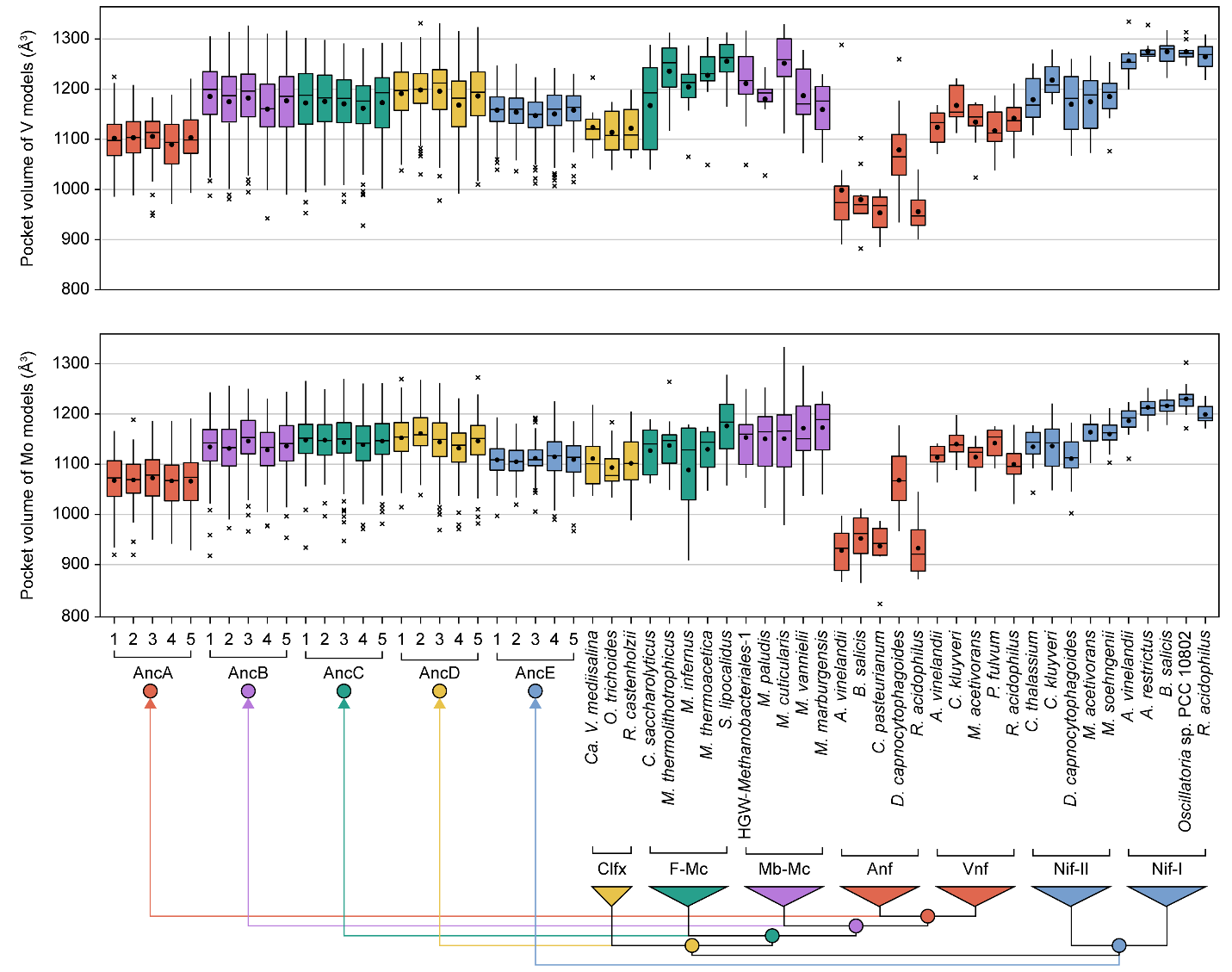

**Figure S4**. Pocket volumes of extant and ancestral nitrogenases modeled with different cofactors. (Top) Volumes for nitrogenases modeled with a FeMo-cofactor and (Bottom) volumes for nitrogenases modeled with a FeV-cofactor. Median values are indicated by bars, mean values by points, the range (excluding outliers) by whiskers, and outliers by crosses. Phylogeny coloring is as follows: Clfx, yellow; F-Mc, green; Mb-Mc, purple; Anf/Vnf, red; Nif-I/-II, blue.

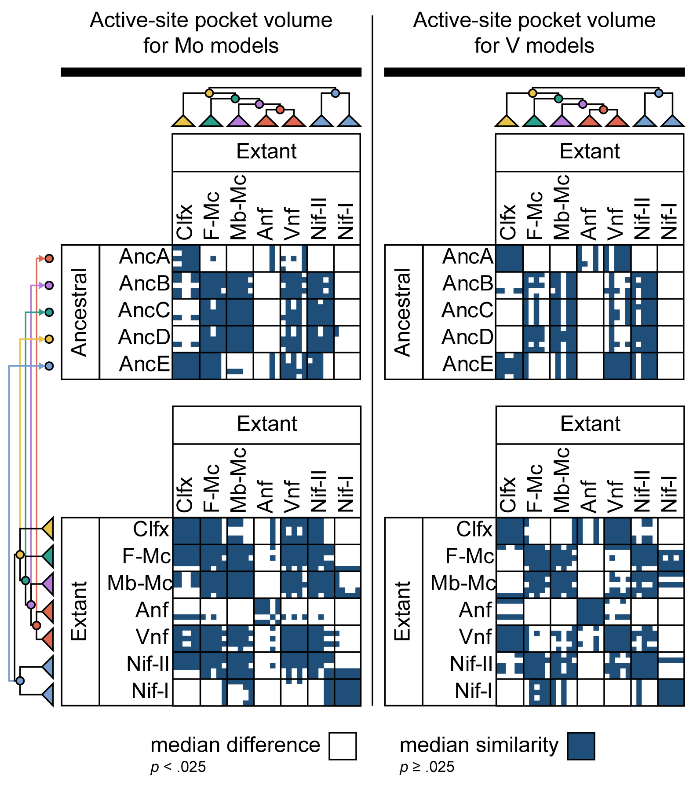

**Figure S5**. Dunn’s test results for pairwise comparisons of modeled nitrogenase pocket volume data, grouped by sequence. Median pocket volume difference for pairwise comparisons is concluded if Dunn’s test null hypothesis is rejected (*p* < .025). Phylogeny coloring is as follows: Clfx, yellow; F-Mc, green; Mb-Mc, purple; Anf/Vnf, red; Nif-I/-II, blue.

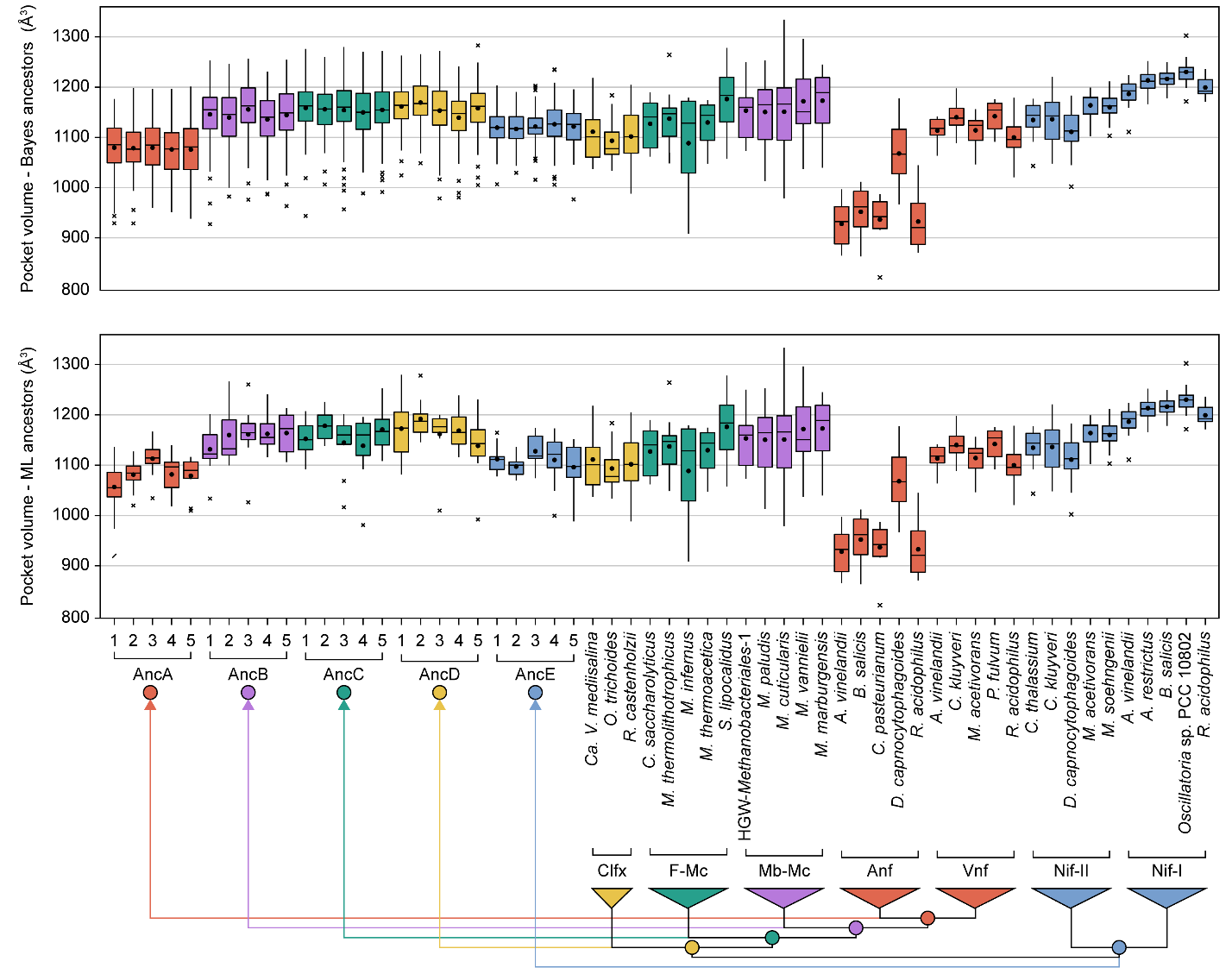

**Figure S6**. Pocket volume of extant and ancestral nitrogenases inferred by different methods. (Top) Volumes for extant and Bayesian-sampled nitrogenases, modeled with a FeMo-cofactor. (Bottom) Volumes for extant and maximum likelihood (ML) nitrogenases, modeled with a FeMo-cofactor. Median values are indicated by bars, mean values by points, the range (excluding outliers) by whiskers, and outliers by crosses. Phylogeny coloring is as follows: Clfx, yellow; F-Mc, green; Mb-Mc, purple; Anf/Vnf, red; Nif-I/-II, blue.
