## Supplemental File 6 for "Reconstructing the Evolutionary History of Nitrogenases: Evidence for Ancestral Molybdenum-Cofactor Utilization"

**Appendix S6: Results of probabilistic model for nitrogenase metal-dependence classification**

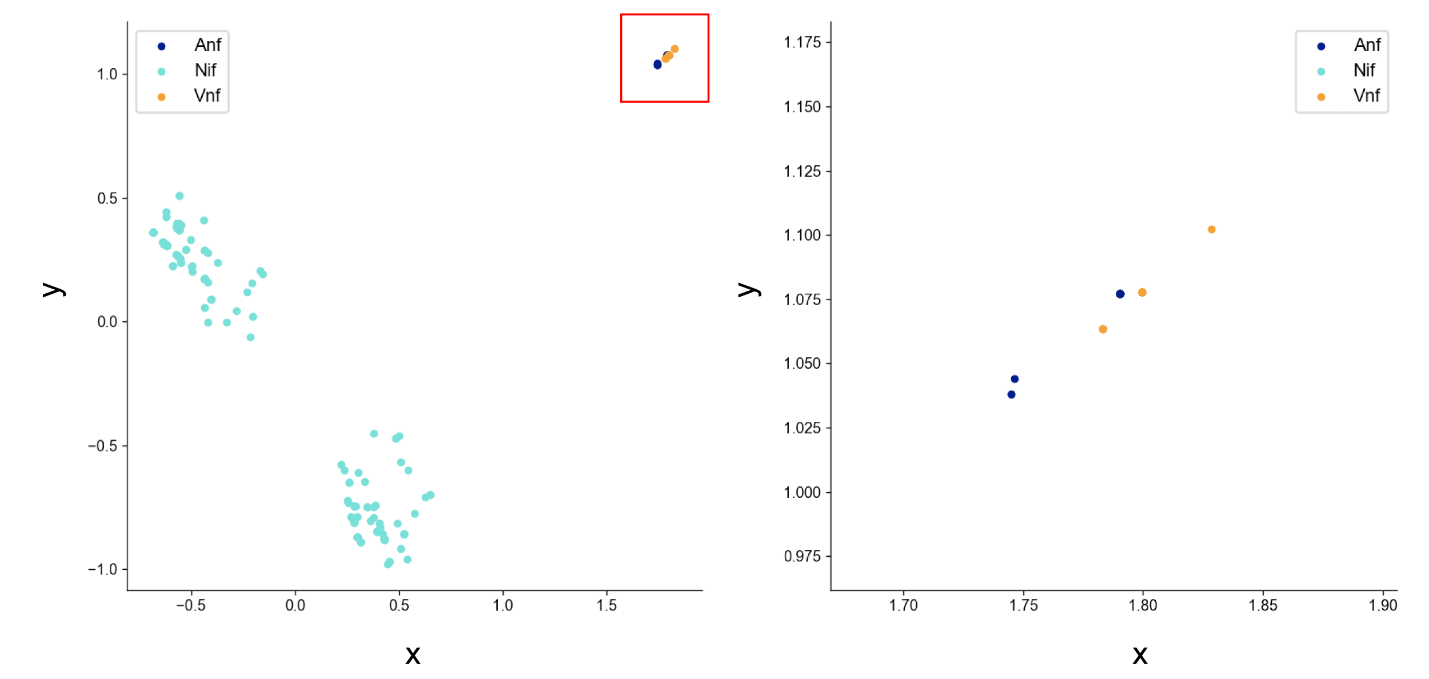

**Figure S6-1.** Principle-component analysis of active-site residue composition of known Nif, Vnf, and Anf extant nitrogenase sequences used to train the classifier. The right panel shows an expanded view of the area labeled by the red square in the left panel.

**Table S6-1.** Nif, Vnf, or Anf classification results for both uncharacterized and maximum-likelihood ancestral nitrogenase sequences. Classification support is reported as the distance of each data sample from the support-vector hyperplane defining the inferred class. A more positive distance value indicates greater support for the inferred class.

| Group | Sequence | Classification | Distance | | |
| --- | --- | --- | --- | --- | --- |
|  |  |  | Nif | Vnf | Anf |
| Clfx | *Ca. V. mediisalina* | Nif | 2.1166 | 0.9572 | -0.0936 |
|  | *O. trichoides* | Nif | 2.1149 | 0.9586 | -0.0925 |
|  | *R. castenholzii* | Nif | 2.1148 | 0.9587 | -0.0924 |
| F-Mc | *C. saccharolyticus* | Nif | 2.1293 | 0.9261 | -0.0863 |
|  | *D. copahuensis* | Nif | 2.1275 | 0.9329 | -0.0896 |
|  | *M. thermolithotrophicus* | Nif | 2.129 | 0.9268 | -0.0864 |
|  | *M. igneus* | Nif | 2.129 | 0.9268 | -0.0864 |
|  | *M. infernus* | Nif | 2.129 | 0.9268 | -0.0864 |
|  | *M. thermoacetica* | Nif | 2.1275 | 0.9329 | -0.0896 |
|  | *S. lipocalidus* | Nif | 2.1293 | 0.9262 | -0.0863 |
|  | *T. phaeum* | Nif | 2.1288 | 0.9269 | -0.0863 |
| Mb-Mc | *M. paludis* | Nif | 2.12 | -0.078 | 0.932 |
|  | *M. cuticularis* | Nif | 2.1311 | -0.0844 | 0.9212 |
|  | *M. vannielii* | Nif | 2.1196 | -0.0778 | 0.9323 |
|  | *M. marburgensis* | Nif | 2.1179 | -0.0766 | 0.9336 |
|  | HGW-*Methanobacteriales*-1 | Nif | 2.1169 | -0.0779 | 0.9366 |
| AncA | AncA-1 | Vnf | 0.9806 | 2.0548 | -0.0396 |
|  | AncA-2 | Vnf | 0.977 | 2.0657 | -0.0488 |
|  | AncA-3 | Vnf | 0.9753 | 2.0676 | -0.0494 |
|  | AncA-4 | Vnf | -0.0884 | 2.1059 | 0.9685 |
|  | AncA-5 | Vnf | 0.9766 | 2.0622 | -0.0444 |
| AncB | AncB-1 | Nif | 2.1373 | 0.9245 | -0.0965 |
|  | AncB-2 | Nif | 2.1381 | 0.9239 | -0.0972 |
|  | AncB-3 | Nif | 2.1372 | 0.9247 | -0.0966 |
|  | AncB-4 | Nif | 2.1207 | 0.9418 | -0.0875 |
|  | AncB-5 | Nif | 2.1384 | 0.9236 | -0.0973 |
| AncC | AncC-1 | Nif | 2.1406 | 0.9177 | -0.0955 |
|  | AncC-2 | Nif | 2.1395 | 0.9188 | -0.0949 |
|  | AncC-3 | Nif | 2.1408 | 0.9173 | -0.0954 |
|  | AncC-4 | Nif | 2.1414 | 0.9179 | -0.0969 |
|  | AncC-5 | Nif | 2.1399 | 0.9186 | -0.0952 |
| AncD | AncD-1 | Nif | 2.1418 | 0.9169 | -0.0967 |
|  | AncD-2 | Nif | 2.141 | 0.9179 | -0.0964 |
|  | AncD-3 | Nif | 2.1423 | 0.9163 | -0.0969 |
|  | AncD-4 | Nif | 2.1426 | 0.917 | -0.0979 |
|  | AncD-5 | Nif | 2.1417 | 0.9175 | -0.0971 |
| AncE | AncE-1 | Nif | 2.1824 | 0.8756 | -0.1267 |
|  | AncE-2 | Nif | 2.182 | 0.8762 | -0.1266 |
|  | AncE-3 | Nif | 2.1807 | 0.877 | -0.125 |
|  | AncE-4 | Nif | 2.1853 | 0.8728 | -0.1295 |
|  | AncE-5 | Nif | 2.1809 | 0.8769 | -0.1251 |
