## Supplementary material for "Reconstructing the Evolutionary History of Nitrogenases: Evidence for Ancestral Molybdenum-Cofactor Utilization": Complete SI List

**Supporting Information List**

Appendix S1: Table of extant nitrogenase protein sequence dataset

- Table S1-1

Appendix S2: Expanded nitrogenase phylogenetic analyses

- Figure S2-1

- Figure S2-2

- Table S2-1

Appendix S3: Expanded nitrogenase pocket volume analyses

- Table S3-1

- Figure S3-1

- Figure S3-2

- Figure S3-3

Appendix S4: Ancestral nitrogenase protein sequence statistical support summary

- Table S4-1

- Figure S4-1

Appendix S5: Table of nitrogenase unique active-site residues

- Table S5-1

Appendix S6: Probabilistic model to classify active-site amino acid probability distributions

- Figure S6-1

- Table S6-1

Files for GitHub:

1) Extant and ancestral nitrogenase protein sequence alignment

- Extant-MLAnc_Align.fasta

2) Bayesian-sampled ancestral nitrogenase protein sequences

- BayesAnc.fasta

3) Homology modelling Modeller scripts

- runModeller_forMo-cofactor,py

- runModeller_forV-cofactor.py

- parseBestModel.py

4) Pocket volume modeling POVME script

- POVME_InputParams.ini

5) Raw pocket volume data

- POVME_Data.csv

6) R scripts for pocket volume plotting/statistical analyses

- Plot_POVMESeq.R

- Stat_POVMESeq.R

7) Python script to build classifier model

- buildClassifierModel.py

8) Output from buildClassifierModel.py

- buildClassifierModelResults.txt
